# Systems-level network modeling deciphers the master regulators of phenotypic plasticity and heterogeneity in melanoma

**DOI:** 10.1101/2021.03.11.434533

**Authors:** Maalavika Pillai, Mohit Kumar Jolly

## Abstract

Phenotypic (i.e. non-genetic) heterogeneity in melanoma drives dedifferentiation, recalcitrance to targeted therapy and immunotherapy, and consequent tumor relapse and metastasis. Various markers or regulators associated with distinct phenotypes in melanoma have been identified, but, how does a network of interactions among these regulators give rise to multiple “attractor” states and phenotypic switching remains elusive. Here, we inferred a network of transcription factors (TFs) that act as master regulators for gene signatures of diverse cell-states in melanoma. Dynamical simulations of this network predicted how this network can settle into different “attractors” (TF expression patterns), suggesting that TF network dynamics drives the emergence of phenotypic heterogeneity. These simulations can recapitulate major phenotypes observed in melanoma and explain de-differentiation trajectory observed upon BRAF inhibition. Our systems-level modeling framework offers a platform to understand trajectories of phenotypic transitions in the landscape of a regulatory TF network and identify novel therapeutic strategies targeting melanoma plasticity.

## Introduction

Melanoma is a highly aggressive cancer arising from melanocytes, the pigment-producing cells of the body. Metastatic melanoma is the deadliest skin malignancy with an abysmal five-year survival rate of 22.5% (Rebecca and Herlyn, 2020). Over 50% of melanomas were found to be driven by BRAF^V600E^ mutation, leading to promising developments in targeted therapy (i.e. BRAF inhibitors - vemurafenib, dabrafenib) over the past decade. However, most of the patients rapidly acquire resistance to these drugs (Sun et al., 2014) and show signs of relapse within 6-8 months of treatment (Sullivan and Flaherty, 2013). This resistance limits cures in metastatic melanoma.

A major hurdle in treating melanoma is the high degree of phenotypic plasticity and heterogeneity. Melanoma comprises cell subpopulations with diverse phenotypes – ‘proliferative’ and ‘invasive’, reminiscent of heterogeneity along the spectrum of epithelial-mesenchymal transition (EMT) seen in carcinomas (Jolly and Celia-Terrassa, 2019). These subpopulations in melanoma show varying drug sensitivities and can interconvert among one another *in vitro* and *in vivo* during metastasis (Hoek et al., 2008; Verfaillie et al., 2015), thus giving rise to ‘moving targets’ that is difficult for a ‘static’ therapy to eliminate. Such reversible cell-state transitions indicate that cellular reprogramming in melanoma is largely driven at transcriptomic and/or epigenetic levels, instead of accumulation of specific DNA mutations. While individual molecules associated with these phenotypes have been identified, a systems-level investigation of the nonlinear dynamics of cell-state transitions in melanoma has not yet been conducted.

MITF, master regulator of melanocytes, is among the most well-studied molecules in melanoma. MITF^high^ cells mark a proliferative phenotype and express melanocytic genes such as MLANA and TYR along with Ki67, while MITF^low^ cells denote an invasive one and express WNT5A and DKK1. Tumors formed by either proliferative or invasive melanoma cell lines contained both MITF^high^ and MITF^low^ cells, indicating bidirectional phenotypic switching between these two (Hoek et al., 2008). An intermediate melanocytic/invasive state expressing NFATC2, SOX6 and ETV4 has also been recently reported (Wouters et al., 2020). Further, in response to BRAF inhibitors, some melanoma cells can exhibit a neural crest stem cell-like (NCSC) state, expressing markers such as NGFR, and proliferating relatively slowly (Fallahi-Sichani et al., 2017; Rambow et al., 2018; Su et al., 2017, 2019; Tsoi et al., 2018). Some of these cells following this de-differentiation trajectory can further progress to an undifferentiated invasive (also called ‘mesenchymal-like’) state, indicating that cells may acquire an NGFR+ state that lies *en route* transitions between proliferative and invasive phenotypes, similar to recent reports about hybrid epithelial/mesenchymal phenotypes (Pastushenko et al., 2018) in EMT. Thus, a gradient of MITF activity (“revised MITF rheostat model”) can define phenotypic heterogeneity in melanoma: the abovementioned four phenotypes (melanocytic/ proliferative, intermediate, NCSC, undifferentiated/invasive) and the two relatively less-characterized ‘hyper-differentiated’ and ‘starved’ ones (**Fig 1A**) (Rambow et al., 2019).

**Fig 1.**
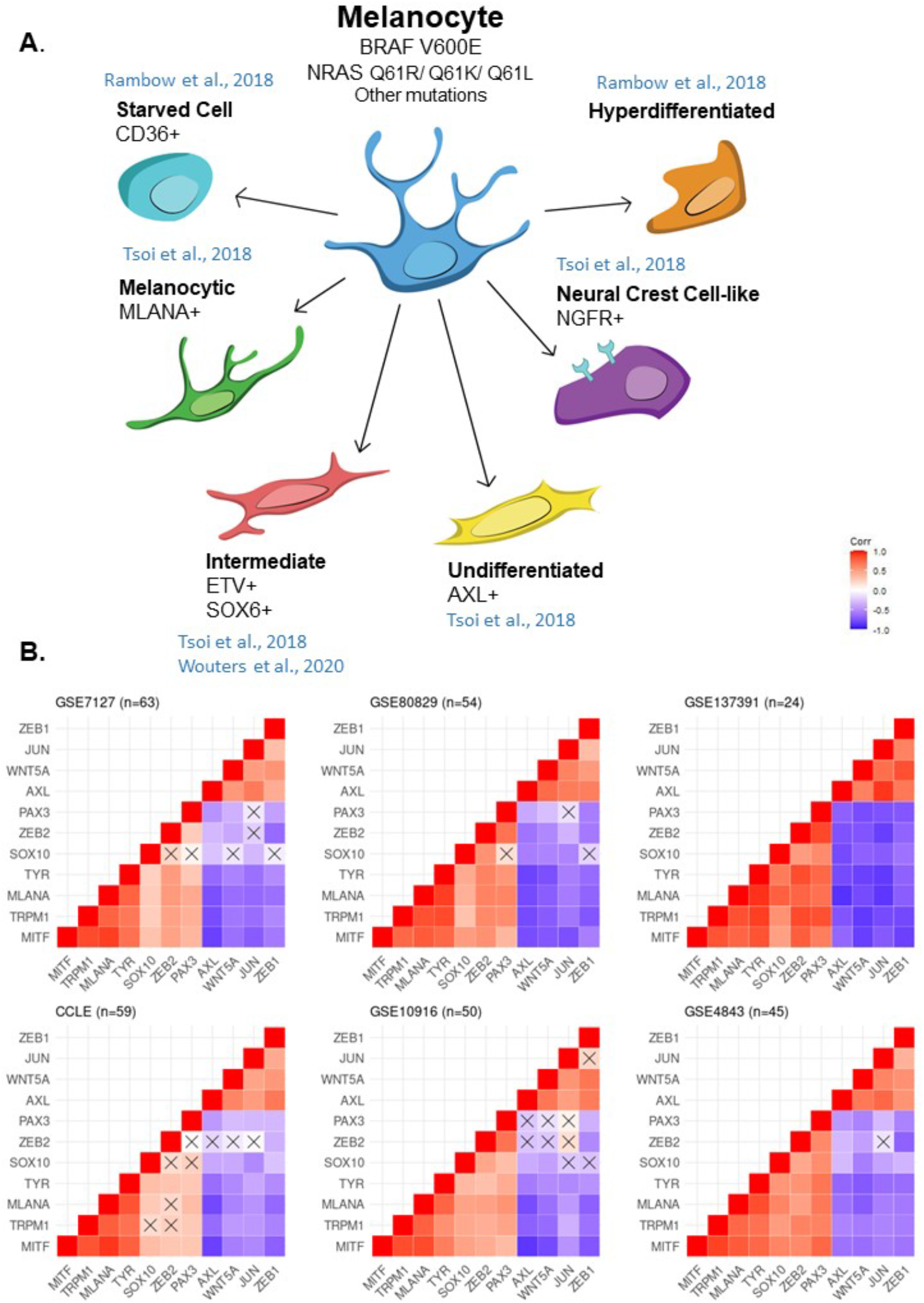
Existence of two distinct groups of genes associated with phenotypic heterogeneity in melanoma. **A.** Experimentally reported phenotypes of melanoma. **B.** Spearman’s correlation of regulators of phenotypic heterogeneity for GSE7127 (n=63), GSE80829 (n=54), GSE137391 (n=24) (left to right, top panel) and CCLE (n= 59 – melanoma cell lines), GSE10916 (n=50) and GSE4843 (n=45) (left to right, bottom panel). Crosses indicate p > 0.05. Colorbar denotes the values of correlation coefficient.

Besides MITF, other reported regulators of phenotypic heterogeneity and/or therapy resistance in melanoma include AXL, c-JUN, BRN2 and PAX3. MITF^low^/AXL^high^ cells can confer resistance to multiple targeted drugs (Müller et al., 2014). Similarly, MITF^low^/c-JUN^high^ cells in mouse and human tumors associate with increased myeloid cell infiltration and inflammation-driven dedifferentiation (Riesenberg et al., 2015). The association of MITF with BRN2 and PAX3 is more complicated: overexpression of BRN2 has been seen to both increase and decrease MITF activity in different cell lines, and a reconciliatory explanation has been postulated involving PAX3 that upon BRAF inhibition, MITF is not activated by PAX3 instead by BRN2 which happens to be a weak activator (Smith et al., 2019). This PAX3/BRN2 rheostat model-based proposed regulation of MITF is also consistent with the identified role of BRN2 in migration of neural crest cells (Fane et al., 2019). Furthermore, ZEB1 and ZEB2 can regulate MITF and their relative levels can associate with a proliferative-invasive switch, as identified by *in vivo* lineage tracing (Vandamme et al., 2020). Put together, these associations and/or regulatory interactions offer important mechanistic insights into the roles of these individual molecules in phenotypic plasticity in melanoma. However, these interactions have not been investigated from a dynamical systems perspective to understand how the emergent properties of a regulatory network involving these players can give rise to multiple “attractor” states in melanoma to explain the diverse set of observations of phenotypic switching.

Here, we integrate correlative metrics among different genes with dynamic modeling of underlying gene regulatory network to explain the emergence of distinct phenotypes in melanoma that can switch among themselves as noted experimentally. Our model simulations can recapitulate the four phenotypes seen in melanoma, identify transcriptional regulators of distinct cell states and phenotypic heterogeneity in melanoma, as well as explain de-differentiation trajectory observed upon BRAF inhibition. This modeling framework offers a platform to predict the diverse dynamic trajectories of phenotypic switching in melanoma and to identify novel therapeutic strategies.

## Materials and methods

### RNA-seq and microarray data

GSE4843 (Hoek et al., 2006) was used for network generation and RACIPE simulations. For other analysis, GSE72056 (Tirosh et al., 2016), GSE134432 (Wouters et al., 2020), GSE112509 (Kunz et al., 2018), GSE137391 (Vivas-García et al., 2020), GSE7127 (Johansson et al., 2007), GSE10916 (Augustine et al., 2010), GSE80829 (Tsoi et al., 2018), Cancer Cell Line Encyclopaedia (CCLE - Broad Institute) (Barretina et al., 2012) and GSE81383 (Gerber et al., 2017) were used. Z-scores were calculated to compare the expression levels of different genes. All analyses have been done using R version 3.6.3, unless mentioned otherwise.

### Correlation plots

To determine the extent of correlation between two genes, we used both Pearson’s correlation coefficient and Spearman’s correlation coefficient. Both methods generate a coefficient ranging between −1 to +1, where +1 indicates a strongly positive correlation and −1 indicates a strongly negative correlation. Pearson’s correlation coefficient is used to determine correlation between linearly related variables whereas Spearman’s correlation coefficient can determine correlation between any monotonically related variable – linear and non- linear. To calculate and plot the correlation matrices *stats* and *ggcorrplot* packages were used.

### K-means clustering

K-means clustering identifies clusters by first, randomly assigning k points in separate clusters and classifying all other points to the clusters by minimizing the sum of squared Euclidean distance of each point from the cluster means. The algorithm calculates cluster centroids and iteratively repeats the assignment of datapoints to the clusters, till no further changes occur. The *kmeans* function from the *stats* package was used to perform k-means clustering on the datasets.

### Selection of optimal number of clusters

Akaike Information Criterion (AIC), Bayesian Information criterion (BIC) and Silhouette scores were used to determine optimal number of clusters to classify the data into. The metrics are calculated as follows:

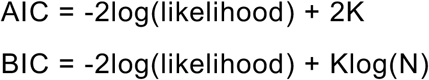

where, K is the number of parameters estimated by each model, and N is the sample size for a given model. For K-means clustering, twice-negative log-likelihood is estimated as the total within cluster sum of squares. To determine the optimal number of clusters, the model with the largest dip in AIC and BIC score was selected. Plots were generated for the difference in score of the n^th^ and (n-1)^th^ model.

### Identification of two phenotypes

K-means clustering was used to generate two clusters of cells based on the top 3000 genes with the highest variance. Each cluster was checked for enrichment of Hoek proliferative, Hoek invasive (Hoek et al., 2008), Verfaillie proliferative and Verfaillie invasive gene sets (Verfaillie et al., 2015) based on signal-to-noise ratio by using the Gene Set Enrichment Analysis (GSEA) software (Subramanian et al., 2005).

### PCA

Principal component analysis (PCA) was used to visualize multidimensional gene expression and simulation data. *factomineR* and *factoextra* packages were used to calculate and visualize the datapoints on the principal components. To determine the correlation between variables and the representation of variables by the principal components, a correlation circle with squared cosines was plotted.

### LDA

Linear discriminant analysis (LDA) is a dimensionality-reduction method that provides linear combinations of variables (discriminant functions) to maximize the separability and differences between multiple classes of data. To identify variables that contribute the most to the separation of two classes of data, we plotted the weights or loading scores of each variable in the first discriminant function. A cut off value of 0.45 was set to select genes that contribute to the separation of the subclusters. The package *MASS* (Venables & Ripley, 2002) was used to calculate the discriminant functions.

### Identification of gene modules for phenotypes

To generate the network underlying the two phenotypes, GSE4843 (Hoek et al., 2006) was subjected to Weighted Gene Correlation Network Analysis (WGCNA) (Langfelder and Horvath, 2008). The topological overlap matrix was constructed based on the Pearson correlation matrix for the gene expression data. A soft threshold power of 4 was identified as the least power that generates a scale free network fit with R^2^ > 0.9. To identify relevant co-expression modules, unsupervised average-linkage hierarchical clustering was used and adaptively pruned at height 0.995. Minimum cluster size was set to 100 to ensure a sufficient size of modules. Merging of modules was repeated at dissimilarity threshold of 0.25 to ensure that similar modules were merged (Tremblay et al., 2019) (**Table S1**).

Modules of interest were identified based on extent of significant differential expression of genes across the two phenotypes based on Bonferroni adjusted p-value (<0.001). Module eigengene values and heatmaps for expression levels of all genes in each module were generated.

### Identification of master regulator network

To identify the master regulators for the differentially expressed genes obtained from WGCNA, we used *geWorkbench* (Floratos et al., 2010). At first, we identified a baseline transcriptional interaction network for the dataset, using ARACNE (Algorithm for the Reconstruction of Accurate Cellular Networks) (Margolin et al., 2006). A p-value of 10^−7^ was set to determine the mutual information threshold and the software was run for 100 bootstraps with data processing inequality set to 0. Fisher’s exact test was used to identify master regulators from a list of candidate master regulators (Lambert et al., 2018). Only those transcription factors (TFs) enriched for in the WGCNA modules with p-value < 0.05 were considered for further analysis. This list was cross validated against CHEA, ENCODE and ARCHS4 databases by using EnrichR (Chen et al., 2013) to identify potential TFs regulating each module. Only those TFs identified by both analyses (ARACNE and EnrichR) were considered as master regulators (**Table S2**).

Interactions between the master regulators was established by manual curation of literature and publicly available databases (**Table S3**). Nodes having no incoming or outgoing edges were removed, to finally arrive at a 17 gene network.

### RACIPE

RAndom CIrcuit PErturbation (RACIPE) (Huang et al., 2017) was used to generate an ensemble of ordinary differential equation (ODE) models. Each model represents a collection of modified Hills equations for each gene, with randomized kinetics parameters sampled from user-defined ranges. Each model is solved for multiple initial conditions, till steady state solutions are obtained. An ensemble of steady state solutions for multiple models is used to represent heterogeneity within a population of cells. For our analysis, we used 10,000 ODE models, each with 100 initial conditions.

### Trajectory analysis

To estimate transcriptional similarity between clusters (i.e. phenotypes) we calculated the Euclidean distance between cluster centres. Euclidean distance (D) between two points is given as the square root of sum of squares of distances and is calculated as follows:

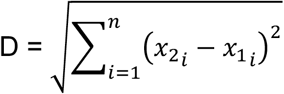

where, n represents number of genes, 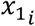 represents value of i^th^ gene for point 1 and 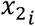 represents value of i^th^ gene for point 2.

### Modelling BRAF/MAPK inhibition

To model the effect of BRAF/MAPK inhibitors on the four phenotypes, we simulated the network using RACIPE with and without MITF knock down (experiment and control cases, respectively), because MITF is a direct target of MAPK and BRAF. To cluster the samples, k-means clustering was used. Both the simulated datasets were projected onto the principal components for the control network and the density distribution of points along PC1 was measured. To determine the Coefficient of Variance (CV), we fit a Gaussian curve to each peak using package *mixtools*. The curve was used to identify the mean (μ) and standard deviation (σ) and calculate CV: CV = σ/μ

### Hartigan’s dip test and bimodality coefficient

To assess the extent of bimodality within genes, we used 2 metric: Hartigan’s dip test and the Bimodality coefficient (BC) (Hartigan and Hartigan, 1985; Pfister et al., 2013). Both measures consider unimodal distributions as their null hypothesis. For Hartigan’s dip test, p-values >0.05 indicate that the distribution is unimodal. To calculate dip test coefficient and p-values, we used the *diptest* package (Maechler, 2016). For BC, values lesser than 0.555 indicate unimodality. To calculate bimodality coefficient, we adapted the *bimodality_coefficient* function from the *modes* package.

## Results

### Two distinct teams of molecules underlying phenotypic heterogeneity

First, we identified a set of genes that regulate and/or correspond to different phenotypes reported in melanoma, based on existing literature. MITF, MLANA, TYR, TRPM1 and other melanocytic genes are upregulated in proliferative cells, while AXL and WNT5A are affiliated with an invasive phenotype (Hoek et al., 2008; Müller et al., 2014; Weeraratna et al., 2002). Further, EMT-inducing transcription factors ZEB1 and ZEB2 can play opposite roles in melanoma, with ZEB2 enabling a more proliferative state while ZEB1 promoting invasion and metastasis (Vandamme et al., 2020). Other molecules such as PAX3, SOX10 and c-JUN regulate phenotypic plasticity too (Riesenberg et al., 2015; Smith et al., 2019). Thus, we calculated pairwise correlations between every two genes, across multiple datasets of melanoma cell lines and human samples.

The correlation matrices revealed an intriguing trend: the invasive phenotype genes ZEB1, JUN, WNT5A and AXL formed one team of players which were all positively correlated with one another (upper red triangle in **Fig 1B**, **S1A**). Similarly, genes corresponding to proliferative phenotype – PAX3, ZEB2, SOX10, TYR, MLANA, TRPM1 and MITF – were positively correlated among one another, forming another such team (lower red triangle in **Fig 1B**, **S1A**). The players of these two teams were found to be negatively correlated with one another (purple rectangle in **Fig 1B**, **S1A**). These results suggest that the proliferative and invasive phenotype genes tend to form two ‘teams’ of mutually opposing players that can govern the emergence of phenotypic plasticity and heterogeneity in melanoma. Such ‘teams’ have been witnessed between EMT-inducing and EMT-inhibiting factors (Jia et al., 2020b), as well as those between master regulators of ‘classic’ and ‘variant’ subtypes in small cell lung cancer (Chauhan et al., 2020), elucidating a potential common design principle for regulatory networks involved in cancer cell plasticity.

### Presence of well-defined proliferative and invasive phenotypes

To examine whether these two ‘teams’ correspond to distinct phenotypes in melanoma in different datasets, we used K-means clustering on top 3000 genes with the highest variance (Tsoi et al., 2018). K-means clustering is used to then identify the optimal number of clusters in these datasets, based on AIC and BIC scores calculated for K=1 to K=15. The highest peak difference in these scores was noticed from K=1 to K=2, indicating that two clusters are optimal (**Fig S1B**).

These two clusters segregate well when projected on their first two principal components (**Fig 2A**). We next checked the enrichment of reported proliferative and invasive signatures in these two clusters across datasets using gene set enrichment analysis (GSEA) (Hoek et al., 2008; Verfaillie et al., 2015). The two clusters obtained for each dataset show enrichment of either the proliferative or invasive signatures reported earlier (**Fig 2B**, **S2A**), thus highlighting that these two clusters can be mapped on to proliferative and invasive phenotypes as initially identified.

**Fig 2.**
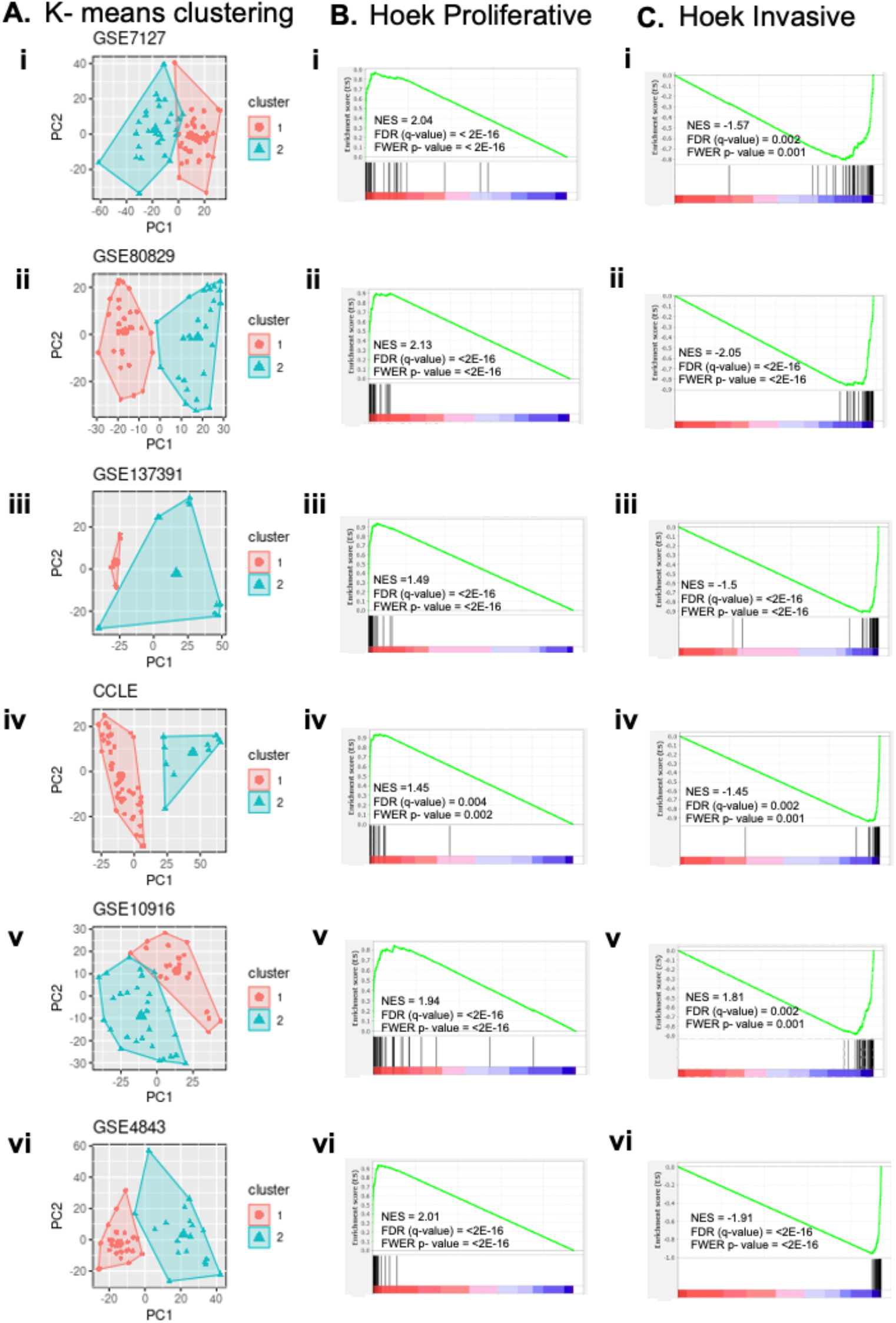
Two distinct classes of cells exist across multiple datasets. **A.** K-means clustering for K=2 yields two distinct clusters along principal component 1 (PC1). GSEA for Hoek proliferative geneset (**B.**) and Hoek invasive geneset (**C.**) confirms that the two clusters correspond to the respective phenotypes for **i**. GSE7127 (n=63) **ii.** GSE80829 (n=53) **iii.** GSE137391 (n=24) **iv.** CCLE (n=59 – melanoma cell lines) **v.** GSE10916 (n=50) **vi.** GSE4843 (n=45)

The presence of these two phenotypic clusters was further confirmed in melanoma samples at bulk (GSE112509) and single-cell (GSE81383) levels (**Fig S2B**). These samples also showed the emergence of two ‘teams’ of players earlier seen for melanoma cell lines (**Fig S2C**). Put together, these results underscore that these two distinct phenotypes: proliferative and invasive can be seen in diverse publicly available melanoma datasets – at bulk and single-cell RNA-seq levels, as well as cell lines and primary/metastatic samples.

### Identifying a master regulator network for phenotypic heterogeneity in melanoma

To identify the genes regulating diverse phenotypes in melanoma, we used an unbiased approach (**Fig 3A**) by first classifying all genes into modules based on correlation among them through WGCNA (Langfelder and Horvath, 2008) (**Fig S3A, i**). These modules comprise co-expressed genes; some of these modules may correspond to specific phenotypes (Udyavar et al., 2017). To calculate the overall expression of each of these modules in a given melanoma tissue sample in GSE4843, we computed corresponding eigengene values (first principal component of all genes within a module). Module eigengene values for salmon and yellow module were significantly different across the proliferative and invasive samples (Bonferroni adjusted p-value <0.001) **(Fig S3A, ii)**. In proliferative samples, eigengene values of yellow module were upregulated and those of salmon module were downregulated (**Fig 3B, i**). Conversely, in invasive samples, eigenvalues of yellow module were downregulated and those of salmon module were upregulated (**Fig 3B, ii**). Thus, these two modules can be considered to correspond to two distinct phenotypes which largely exhibit mutually exclusive gene expression patterns (**Fig S3A, iii)**.

**Fig 3.**
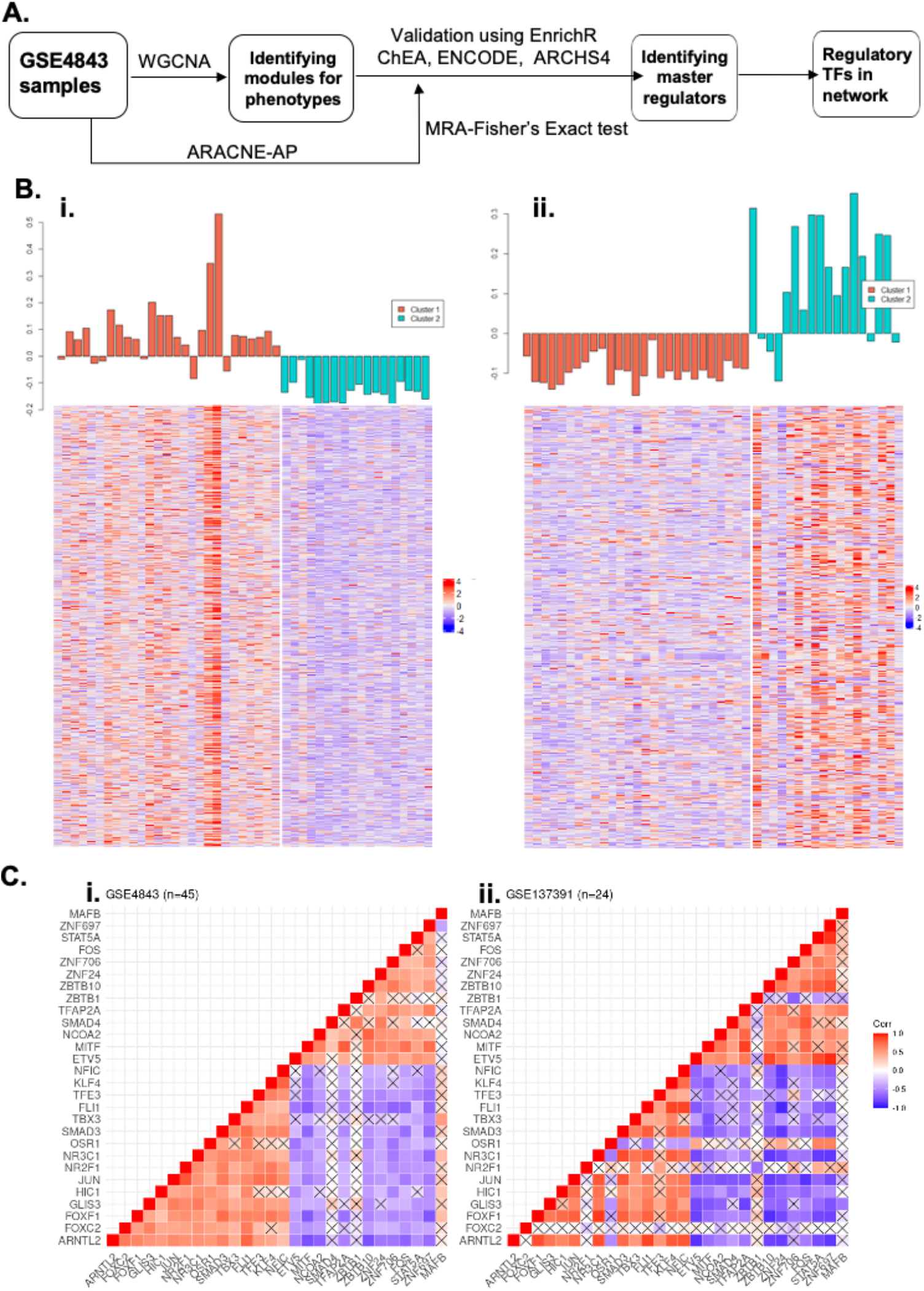
Identification of master regulators driving phenotypic heterogeneity in melanoma. **A.** Pipeline followed to identify nodes for underlying TF regulatory network. **B.** Modules identified from WGCNA for GSE4843: **i.** Yellow module **ii.** Salmon module. Module eigengene values are shown for each sample (top) and the corresponding expression level heatmap of genes in each module (bottom) for proliferative (orange) and invasive (cyan) samples are given for GSE4843. **C.** Spearman’s correlation matrix among pairs of master regulators in: **i.** GSE4843 and **ii.** GSE137391. Crosses indicate p > 0.05. Colorbar denotes correlation coefficient.

Next, we sought to decipher the master regulators of these two modules of interest, using two orthogonal and unbiased approaches. First, we use EnrichR (Chen et al., 2013) which scans publicly available databases to identify potential regulators of genes comprising our modules of interest. Second, we infer a backbone regulatory network using ARACNE (Margolin et al., 2006) on GSE4843 for a set of transcriptional regulators. Nodes in this network were selected based on Fishers’ exact test to identify the master regulators corresponding to each module (Lambert et al., 2018). Finally, we found common regulators identified by these two complementary approaches. The master regulators identified formed two mutually inhibiting groups or ‘teams’, similar to the interactions seen in known regulators of phenotypic heterogeneity, across different *in vitro* and tumor sample datasets (**Fig 3C**, **S3B**). The network obtained via ARACNE is undirected in nature. Thus we arrived at a master regulatory network for melanoma by collating information on directed links in the network (**Fig 4A**) through manual curation of the above-mentioned regulators by using public databases (ChEA, ENCODE) and examining relevant experimental literature (**Table S1**).

**Fig 4.**
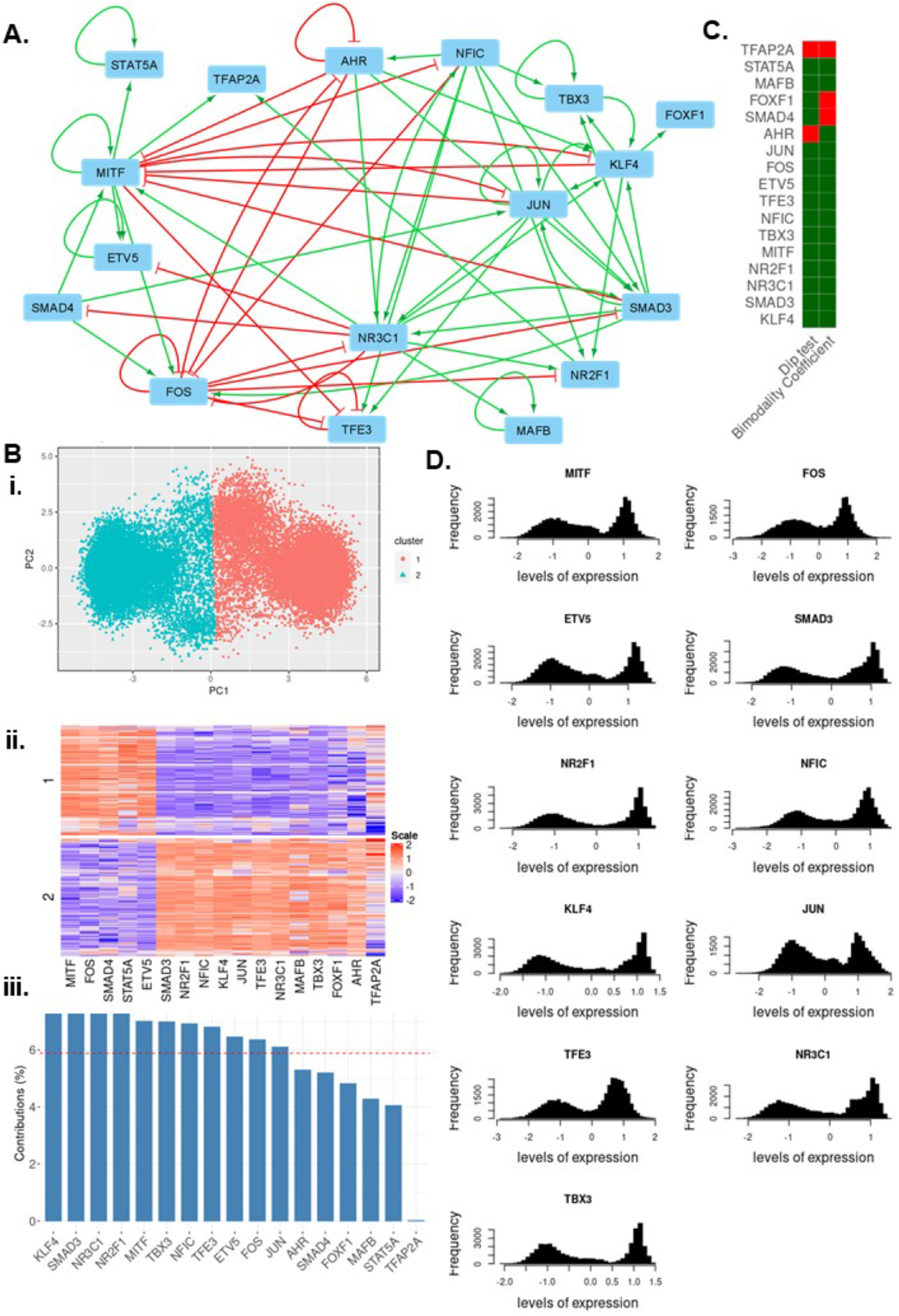
Dynamic simulations recapitulate phenotypic heterogeneity in melanoma. **A.** Interaction network identified for master regulators. **B. i.** PCA plot and **ii.** Heatmap for simulated data forming two distinct clusters **iii.** Percentage contribution of each gene to PC1. The red line indicates the value for uniform contribution (= 1/17) of all genes considered (n=17).**C.** Bimodality test using Bimodality coefficient (BC) (Green boxes indicate BC > 0.555 and red boxes indicate BC < 0.555) and Hartigan’s dip test (Green boxes indicate p < 0.05 and red boxes indicate p > 0.05). **D.** Histograms for individual gene expression distribution in the simulated dataset.

### Emergent network dynamics recapitulate phenotypic heterogeneity in melanoma

Next, we simulated the dynamics of the 17-node regulatory network identified (**Fig 4A**) using an ensemble of kinetic parameter sets to capture the cell-to-cell variability in gene expression levels and/or bio-chemical reaction rates, through the RACIPE (Random Circuit Perturbation) method (Huang et al., 2017). The steady state values of expression levels obtained via RACIPE (further referred to as the simulated dataset) for each gene captures the possible expression levels of these genes within a cell. The simulated dataset had optimal clustering for K=2, based on silhouette width estimation for K-means clustering (K=2 to 10) (**Fig S4A, i**). The two clusters were spatially separated along the first principal component (PC1) (**Fig 4B, i**) and exhibit differential expression of several genes in the network (**Fig 4B, ii**). To identify relevant genes involved in this classification, we calculated the contribution of each gene to PC1 (**Fig 4B, iii**) and the extent of representation of each gene by PC1, based on their squared cosines (**Fig S4A, ii**). Based on both of these metrics, MITF, FOS, KLF4, NFIC, TBX3, NR2F1, SMAD3, ETV5, JUN, TFE3 and NR3C1 were considered to be important for segregation of the two clusters. The distributions for these genes exhibit bimodality (**Fig 4C,D)**, further supporting their differential expression in two clusters.

Several poorly contributing genes (AHR, SMAD4, FOXF1, MAFB, STAT5A, TFAP2A – **Fig 4B, iii**) do not exhibit significant bimodality based on Hartigan’s dip test and Bimodality Coefficient (Hartigan and Hartigan, 1985; Pfister et al., 2013) (**Fig 4C**, **S4B**) in the simulated data. Therefore, these genes are less likely to be able to distinguish between the two clusters.

To confirm whether these two clusters correspond to the two phenotypes (proliferative, invasive), we compared the simulated dataset with multiple experimental datasets for the expression of 11 genes relevant to segregation along PC1 (MITF, FOS, KLF4, NFIC, TBX3, NR2F1, SMAD3, ETV5, JUN, TFE3 and NR3C1). Proliferative and invasive samples (as identified earlier in **Fig 2A** by GSEA of respective signatures) exhibit differential expression of these 11 genes, as predicted by the simulated data (**Fig 5A**, **S4C**). The samples are also spatially separated along the first two principal components for these 11 genes, across diverse *in vitro* and tumor sample datasets (**Fig 5B**). Additionally, correlation plots for these datasets and the simulated dataset show similar patterns, where these 11 genes seem to split into forming two teams of 8 genes and 3 genes (**Fig 5C**, **S4D**), similar to our earlier observations (**Fig 1B**). One of these teams includes MITF, and the other includes JUN, thus reminiscent of antagonism between them and correspondingly opposite phenotypes associated with these molecules. Together, this analysis confirmed that dynamic simulations for the master TF regulatory network we inferred can recapitulate the existence of the proliferative and invasive phenotypes.

**Fig 5.**
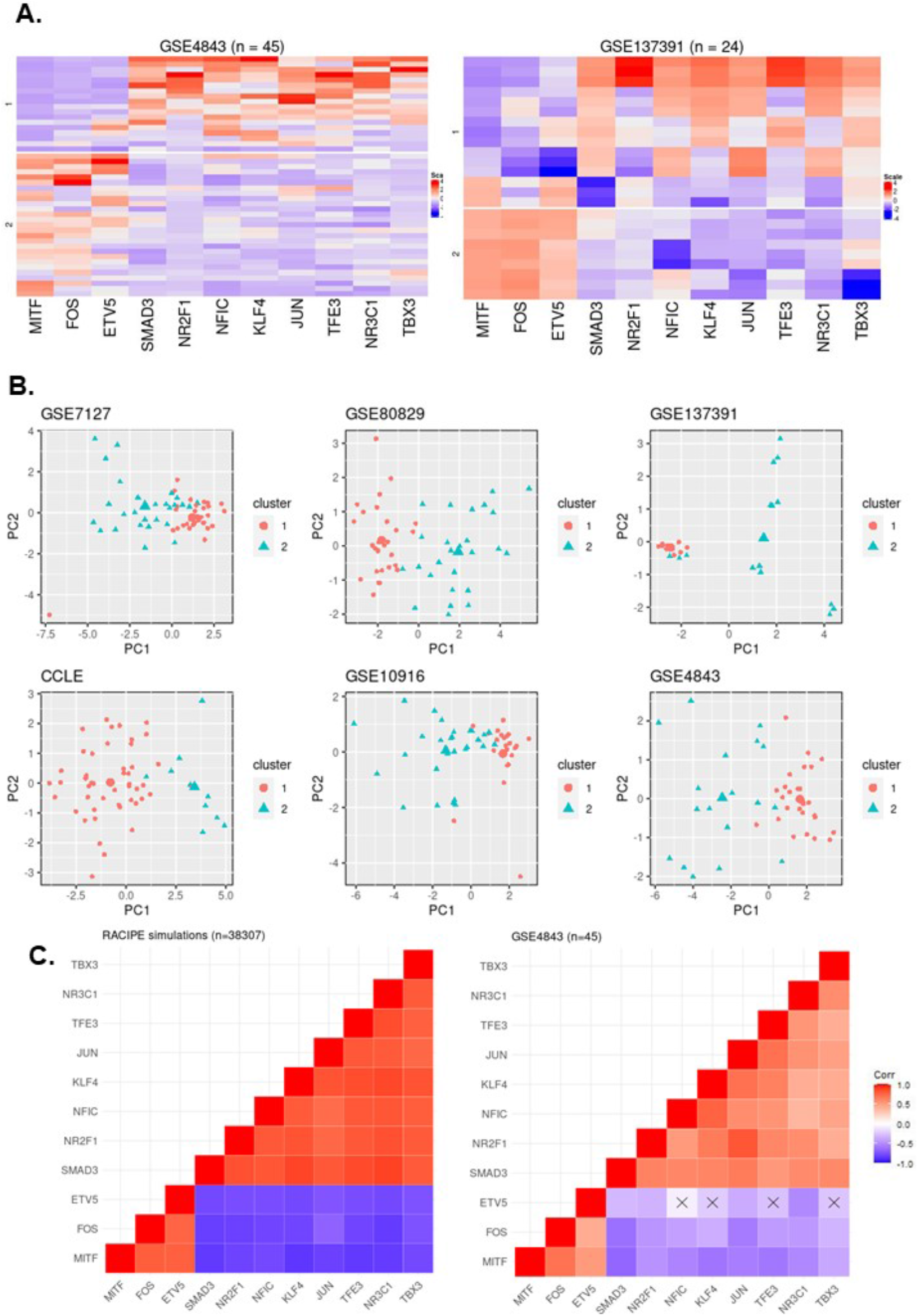
A subset of master regulators explain the existence of two phenotypes. **A.** Differential expression of subset of master regulators (n = 11) in proliferative and invasive samples in GSE4843 and GSE137391. **B.** Projection of proliferative and invasive clusters on the principal components for expression levels of these 11 master regulators. **C.** Spearman’s Correlation coefficient matrix for the 11 master regulators in **i.** RACIPE simulated dataset **ii.** GSE4843. Crosses indicate p > 0.05. Colorbar denotes correlation coefficient.

### Relative proximity and transition paths among four co-existing phenotypes in melanoma

Because the mean silhouette scores for K=3 and K=4 suggested their ability to offer efficient clustering as well (**Fig S4A, i**), and previous reports pinpoint four possible phenotypes in melanoma (Tsoi et al., 2018), we looked at the possibility of four clusters corresponding to the four phenotypes. In the simulated data, these four clusters form distinct groups along the first two principal components (**Fig 6A, i**). Interestingly, the four clusters appear to be further classification of the two clusters previously observed (**Fig 6A, ii** **and** **Fig 4B, ii**), similar to the classification observed in experimental data (Tsoi et al., 2018). To identify genes that distinguish between the proliferative and invasive sub-clusters, we used Linear Discriminant Analysis (LDA). LDA identifies a linear combination of genes that maximizes separability of clusters, thus, the loading scores of each gene were analysed to identify genes that contribute the most. A cut-off of 0.6 was set to classify a gene as a discriminant. MITF was identified as a discriminant between the two proliferative sub-clusters (**Fig S5A, i**) and TBX3, NR2F1, KLF4 and FOXF1 as discriminants for the two invasive sub-clusters (**Fig S5A, ii**). To check if these four clusters map onto the four phenotypes observed, we examined the expression of these determinant genes in experimental datasets. To classify a given dataset into these four phenotypes, we used K-means clustering on the basis of previously defined markers of the 4 phenotype – *MLANA* for the melanocytic phenotype, *ETV4* for intermediate phenotype, *AXL* for undifferentiated phenotype and *NGFR* for NCSC (**Fig 6B**, **S5B**). Differential expression of discriminant genes was observed at single cell resolution in melanoma cell lines (GSE134432) as well as in tissue culture samples from tumours (GSE4843) (**Fig 6C**). This analysis suggests that the four phenotypes observed in melanoma can exist at an individual cell level as well, thus facilitating heterogeneity at a bulk level.

**Fig 6.**
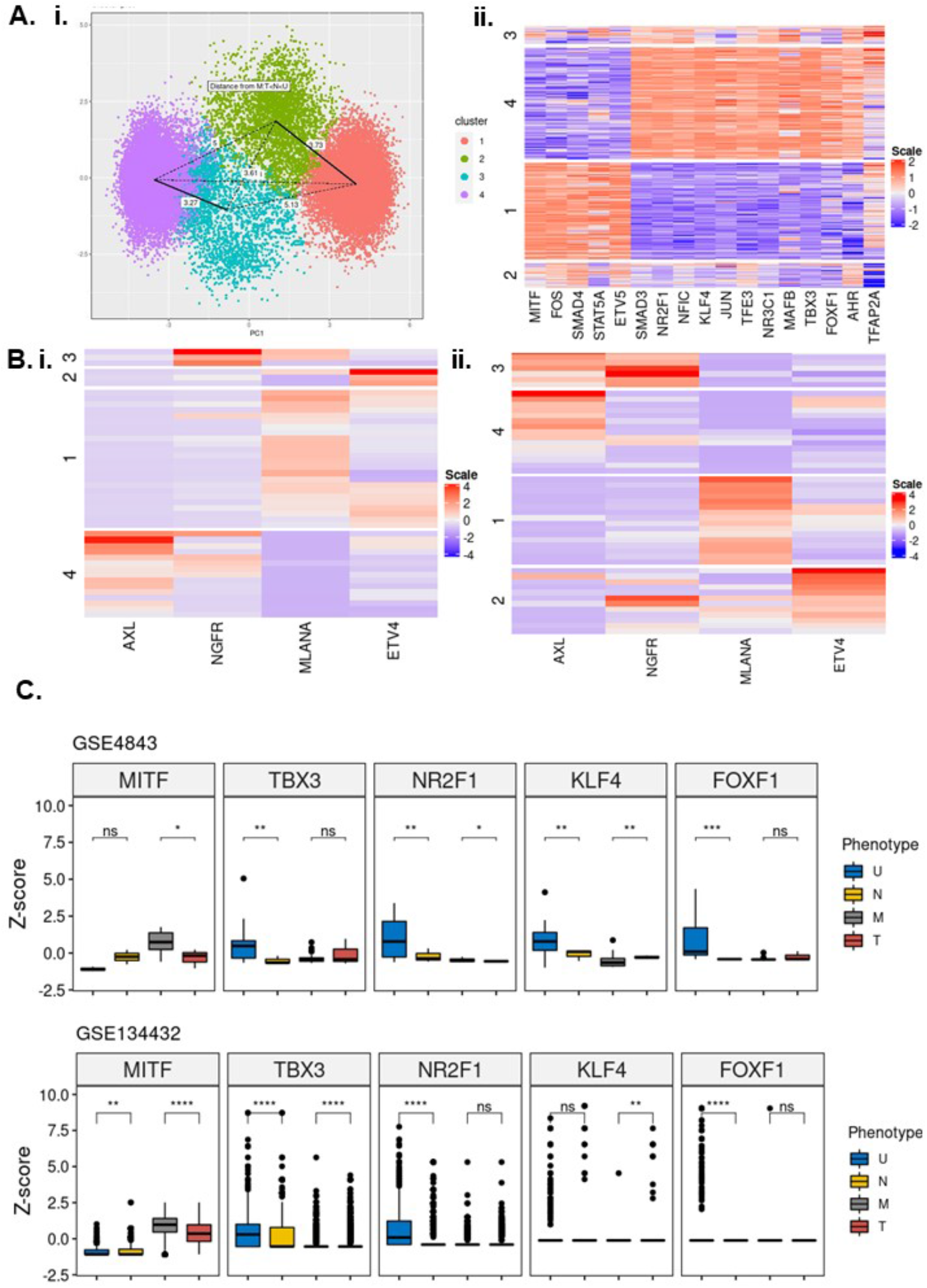
Master regulator network explains the existence of proliferative and invasive subpopulations and dedifferentiation trajectory followed during BRAF inhibition. **A**. **i**. Heatmap and **ii**. PCA plot for four clusters identified in simulated data **B**. Heatmap of marker genes expression levels for the 4 phenotypes (AXL, NGFR, MLANA and ETV4). **i**. GSE4843 **ii**. GSE10916. **C**. Z-scores for discriminant genes can distinguish phenotypes within the proliferative subclusters (Melanocytic – M and Transitory -T) and invasive subclusters (Undifferentiated -U and NCSC -N) in GSE4843 and GSE134432. (Significance is represented based p-value for Student’s t-test, * for p< 0.1, ** for p < 0.01, *** for p < 0.001 and **** for p < 0.0001).

Interestingly, we observed that the proximity between clusters for the simulated dataset reflected features of the de-differentiation trajectory followed during drug treatment (Su *et al.*, 2017; Tsoi *et al.*, 2018). Based on the projection of expression data on the PC axes (**Fig 6A, i)**, the melanocytic cluster is found to be closest to the transitory cluster, and the undifferentiated invasive cluster is closest to the NCSC one, suggesting that during their transition to undifferentiated phenotype, melanocytic cells pass through intermediate phenotypes that are transcriptionally similar. To confirm this prediction about relative positioning of the phenotypes in the experimental datasets, we measured the shortest Euclidean distance between the clusters, based on expression levels of master regulators (**Fig S5C**). Validating the predictions made by the model, the intermediate phenotypes (transitory and NCSC) are found to be closest to the melanocytic and undifferentiated phenotypes which explains the path followed in the de-differentiation trajectory (from melanocytic to transitory to NCSC to undifferentiated). Thus, the cell-fate landscape formed by the emergent dynamics of master TF network can not only explain the existence of four phenotypes earlier reported in melanoma, but also identify most likely transition paths or trajectories that cells follow while transitioning among these states.

### Network-based modelling explains population dynamics during BRAF inhibition

Resistance to MAPK/BRAF inhibitors is accompanied by an increase in invasive cell population (Hoek et al., 2008; Su et al., 2017, 2019). We wanted to check if our network could explain such phenotypic switching in the presence of drug treatment. One of the downstream targets of MAPK and BRAF is MITF. Hence, to represent drug treatment, we simulated the network under MITF knockdown (KD) conditions using RACIPE. Interestingly, we observed a shift in the density of steady-state solutions (i.e. phenotypic composition) from proliferative towards invasive phenotypes (**Fig S6A**). This shift is caused by a significant decrease in the frequency of cells belonging to melanocytic cluster (**Fig 7A**). Although there is no noticeable change in proportion of undifferentiated phenotype, we see a significant increase in the frequencies of the transitory and NCSC cell phenotypes. This change is also accompanied by an increase in the coefficient of variance for the distributions corresponding to the proliferative and invasive phenotypes, suggesting higher phenotypic heterogeneity in response to the drug, and also an increased dominance of these intermediate phenotypes in the overall cell population composition (**Fig 7B**). Similar trends have also been observed experimentally, where short term MAPK inhibition leads to a large increase in Starved Melanoma Cell (SMC, a possible form of the transitory phenotype) and NCSCs (Rambow et al., 2018).

**Fig 7.**
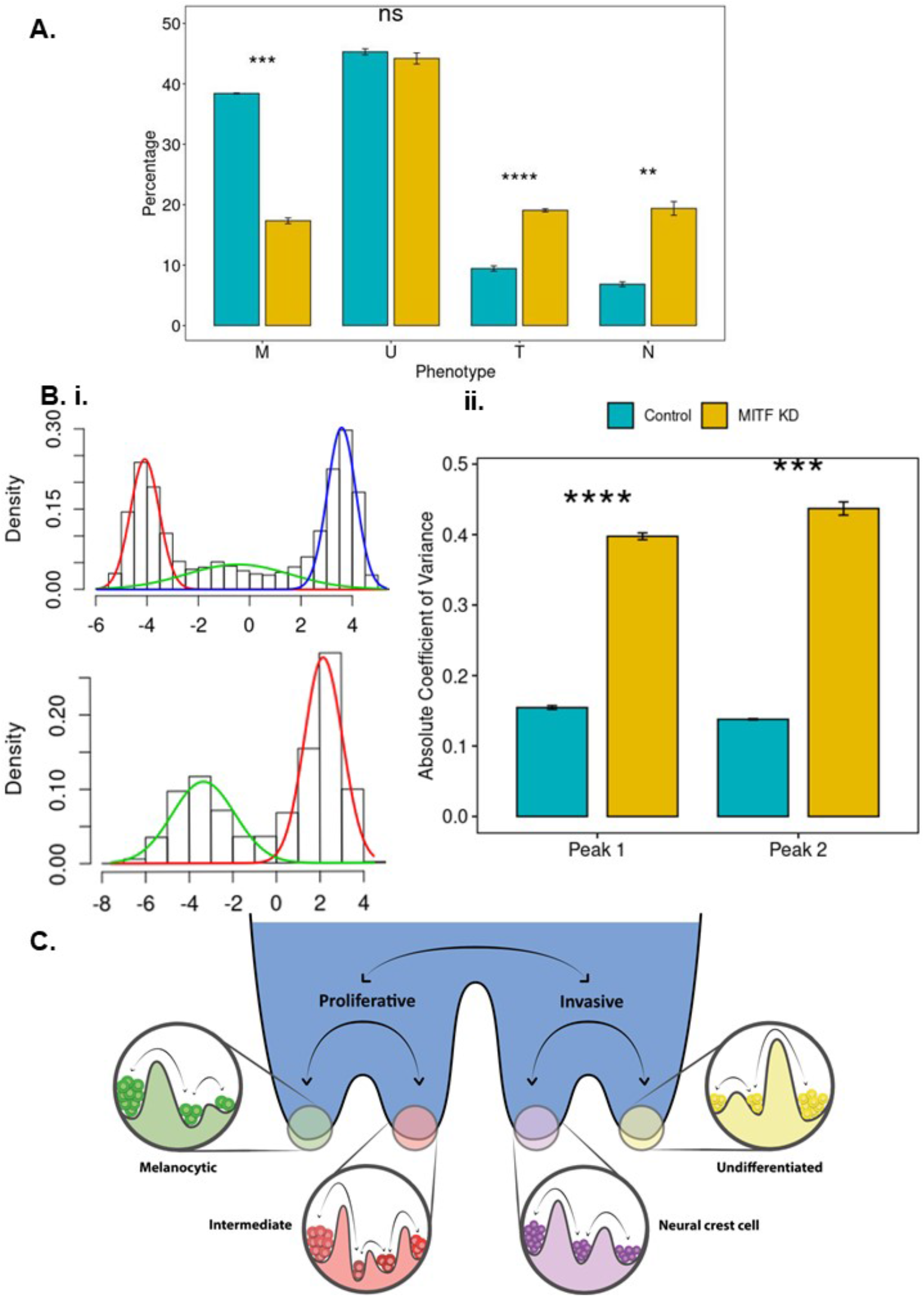
MITF knockdown explains phenotype switching caused by BRAFi and MAPKi. **A.** Barplot representing percentage of solutions representing each phenotype **B.** Density of datapoints form a multimodal distribution along PC1 in Control (Top) and MITF-KD (bottom) networks [left]. MITF-KD increases coefficient of variance of the two gaussian distributions corresponding two the proliferative and invasive phenotypes [right]. (n=3, error bars represent SD, Significance is represented based p-value for Student’s t-test, * for p< 0.05, ** for p < 0.01, *** for p < 0.001 and **** for p < 0.0001). **C.** Cellular plasticity gives rise to phenotypic heterogeneity in melanoma.

To identify whether knocking down any of the other genes in our master TF network can also drive phenotypic switching, we performed *in silico* knockdowns of each gene. Our results showed that knockdown of none of the other genes was as potent as that of MITF in terms of inducing phenotypic switching, suggesting that this effect is unique to MITF knock down (**Fig S6B**). Such a drastic effect observed upon perturbing MITF is very likely a function of the underlying network topology that confers the status of ‘master regulator’ to MITF in defining the cell-state landscape.

## Discussion

Phenotypic heterogeneity in melanoma has been associated with drug sensitivity, immunotherapy resistance and dedifferentiation of melanoma cells (Reinhardt et al., 2017; Su et al., 2017; Tsoi et al., 2018). Here, we identify a transcription factor (TF) regulatory network whose emergent dynamics can settle into multiple “attractors” states, each corresponding to a specific phenotype. The concept of “attractors” is embedded in the mathematical realization of the Waddington’s epigenetic landscape, a powerful approach to understand the dynamics of cellular differentiation in development and increasingly in cancer too (Li et al., 2016; Su et al., 2017; Zhou and Huang, 2011). The four phenotypes seen for melanoma is a refined classification of two broader states: proliferative and invasive, similar to the classification system used previously (Tsoi et al., 2018). Thus, our model proposes the possibility of two “macro-states” corresponding to the proliferative and invasive phenotypes, that comprise “micro-states” corresponding to the sub-clusters (**Fig 7C**). These microstates can further be constituted of an ensemble of microstates; for instance, the hyperdifferentiated cells may be thought of as a microstate for the melanocytic phenotype, and starved melanoma cell (SMC) for the intermediate phenotype (Rambow et al., 2019). Transitions among microstates have lower barriers and are postulated to occur relative easily as compared to transition between macro-states, which might need a stronger “driving force” such as drug treatment (i.e. BRAF inhibition) or genomic silencing. A similar model has been proposed in the context of EMT (Goetz et al., 2020), where such “micro-states” may facilitate cellular adaptation.

Several parallels can be drawn between phenotypic switching in melanoma and EMT in epithelial cancers. First, the invasive phenotype has upregulation of several mesenchymal genes and EMT-regulators such as ZEB1, WNT5A, CDH2 (Su et al., 2017; Vandamme et al., 2020), as well as that of matrix metalloproteases (MMPs) that can change the extra-cellular matrix (ECM) stiffness (Arozarena and Wellbrock, 2017), as typically observed in EMT (Deng et al., 2020; Wei et al., 2015). However, ZEB1 and ZEB2, both EMT-inducing factors, promote opposite phenotypes in melanoma (Denecker et al., 2014). Other EMT regulators such as TWIST and AP-1 (Feldker et al., 2020) are also reported to drive phenotypic plasticity in melanoma (Caramel et al., 2013; Verfaillie et al., 2015). Second, heterogeneity along the EMT axis seen in circulating tumor cells (CTCs) (Bocci et al., 2021; Yu et al., 2013) is recapitulated in melanoma CTCs too (Aya-Bonilla et al., 2020). Third, cooperation among phenotypes with varying EMT status has been witnessed during metastasis and tumor formation (Jolly and Celia-Terrassa, 2019), reminiscent of recent observations in melanoma (Rowling et al., 2020). Further investigations into the dynamics of phenotypic transitions can elucidate whether melanoma cells exhibit hysteresis (Karacosta et al., 2019) and/or spontaneous/stochastic state switching (Tripathi et al., 2020) that characterize EMT.

Our study characterizes phenotypes based on dynamics of TF regulatory network. However, previous studies have also identified translational, epigenetic and metabolic features of distinct phenotypes, suggesting multiple interconnected levels of regulation of phenotypic plasticity in melanoma (Bettum et al., 2015; Falletta et al., 2017; Hugo et al., 2015; Lionetti et al., 2020). Variations in chromatin accessibility, DNA and histone modifications can be mapped to transcriptional heterogeneity seen in melanoma (Hugo et al., 2015; Verfaillie et al., 2015). Two regulatory networks corresponding to proliferative and invasive phenotypes have been identified, with SOX10 and MITF acting as key regulators of the proliferative network and TEAD and AP-1 as regulators of the invasive one. How epigenetic silencing enables reversible or irreversible transitions (Jia et al., 2019) in melanoma, remains to be yet investigated. Besides diverse epigenetic signatures, the proliferative and invasive phenotypes have been characterized to have distinct metabolic profiles too. A switch from proliferative to invasive phenotypes is supported by metabolic reprogramming from oxidative phosphorylation (OXPHOS) to glycolysis mediated by MITF target gene, PGC1α which promotes OXPHOS. Lowering MITF and PGC1α can shift cells towards glycolysis (Vazquez et al., 2013). A recent theoretical model predicted the possibility of four metabolic profiles - OXPHOS-high/glycolysis-low, OXPHOS-low/ glycolysis-high, OXPHOS-low/glycolysis-low, and OXPHOS high/glycolysis-high (Jia et al., 2020a). The first two profiles can be mapped onto the proliferative and invasive phenotypes, based on previous experimental evidence. The ability of intermediate phenotype to capture features of proliferative and invasive cells raises the possibility of its ability to exhibit both OXPHOS and glycolysis. The low/low phenotype, has been identified to map onto an idling population of cells, with no net growth. NCSCs, considered to be slow-growing and to maintain minimal residual disease (Liguoro et al., 2020), may possibly represent the ‘idling’ population of melanoma cells (Paudel et al., 2018) exhibiting both low glycolysis and OXPHOS.

The physiological relevance of phenotypic switching is observed in drug resistance in melanoma. The more invasive phenotypes, NCSC and undifferentiated cells, are observed to emerge along differing time scales upon drug exposure. While shorter duration of BRAF and RAF/MEK inhibition gives rise to NCSC phenotype (NGFR+ cells), prolonged exposure facilitates undifferentiated cells (Fallahi-Sichani et al., 2017; Su et al., 2017). How does adaptation at different time scales contribute to emergence and maintenance of drug-tolerant persisters (Ahmed and Haass, 2018; Schuh et al., 2020; Shaffer et al., 2017), and the role of reversible cellular reprogramming (Karki et al., 2021; Roesch et al., 2010) in enabling such persisters requires further careful investigation. Understanding the dynamics and trajectories of phenotypic switching in these scenarios can help guide better combinatorial (Boshuizen et al., 2018; Luo et al., 2018) and/or sequential (Goldman et al., 2015) therapies that can target vulnerabilities of diverse phenotypes. Our simulations were able to recapitulate the phenomenon of phenotypic switching observed upon BRAF inhibition, thus offering our mechanism-based model as a possible platform to identify potent interventions.

Overall, we have identified a master TF regulatory network that explains the (co-)existence of diverse phenotypes observed in melanoma cell lines and tumours, and the phenomenon of phenotypic switching, which is crucial in melanoma development, progression and metastasis. Our model depicting different “attractor” states also provides a framework to develop future therapeutic strategies, by identifying the key drivers of distinct drug-resistant subpopulations. By targeting the master regulators of given phenotype(s), we can “push” the cells towards more drug-sensitive phenotypes, thereby improving the outcomes of conventional therapy options in use.

## Author contributions

MKJ designed and supervised research. MP performed research and analysed data. Both authors contributed to writing the manuscript.

## Code availability

All codes used in the manuscript are available at: https://github.com/csbBSSE/Melanoma

## Acknowledgement

This work was supported by Ramanujan Fellowship awarded to MKJ by Science and Engineering Research Board (SERB), Department of Science and Technology (DST), Government of India (SB/S2/RJN-049/2018) and by Infosys Young Investigator award to MKJ supported by Infosys Foundation, Bangalore. MP is supported by KVPY fellowship (DST). Atchuta Srinivas Duddu is acknowledged for artwork (**Fig 1A**, **7C**).

---

**Fig S1.**
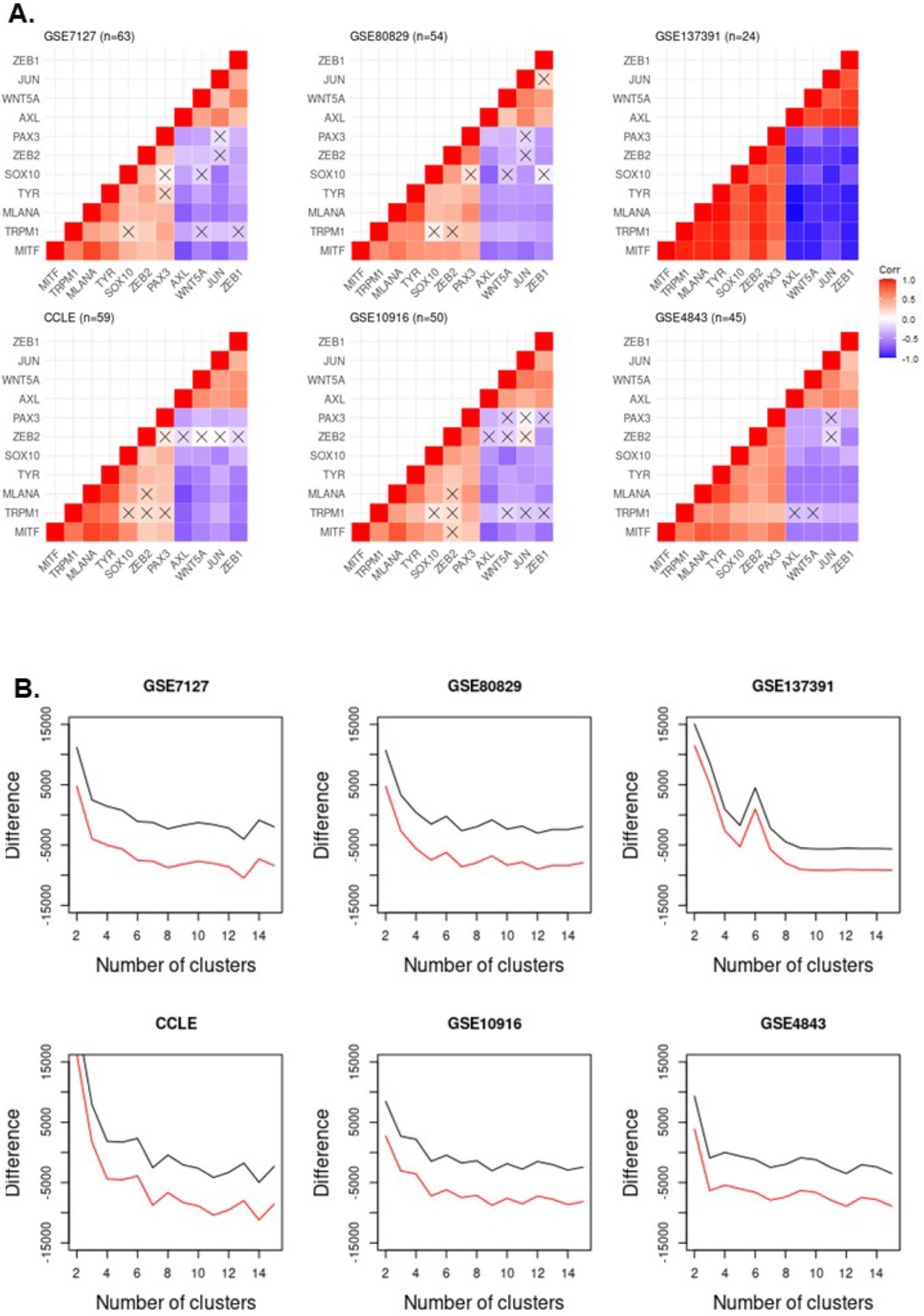
**A.** Pearson’s Correlation of regulators of phenotypic heterogeneity for GSE7127, GSE80829, GSE137391 (left to right, top panel) and CCLE, GSE10916, GSE4843 (left to right, bottom panel). Crosses indicate p > 0.05. Colorbar denotes correlation coefficient. **B.** Difference between n^th^ and (n-1)^th^ AIC (red line) and BIC (black line) scores for 2-15 clusters for GSE7127, GSE80829, GSE137391 (left to right, top panel) and CCLE, GSE10916, GSE4843 (left to right, bottom panel)

**Fig S2.**
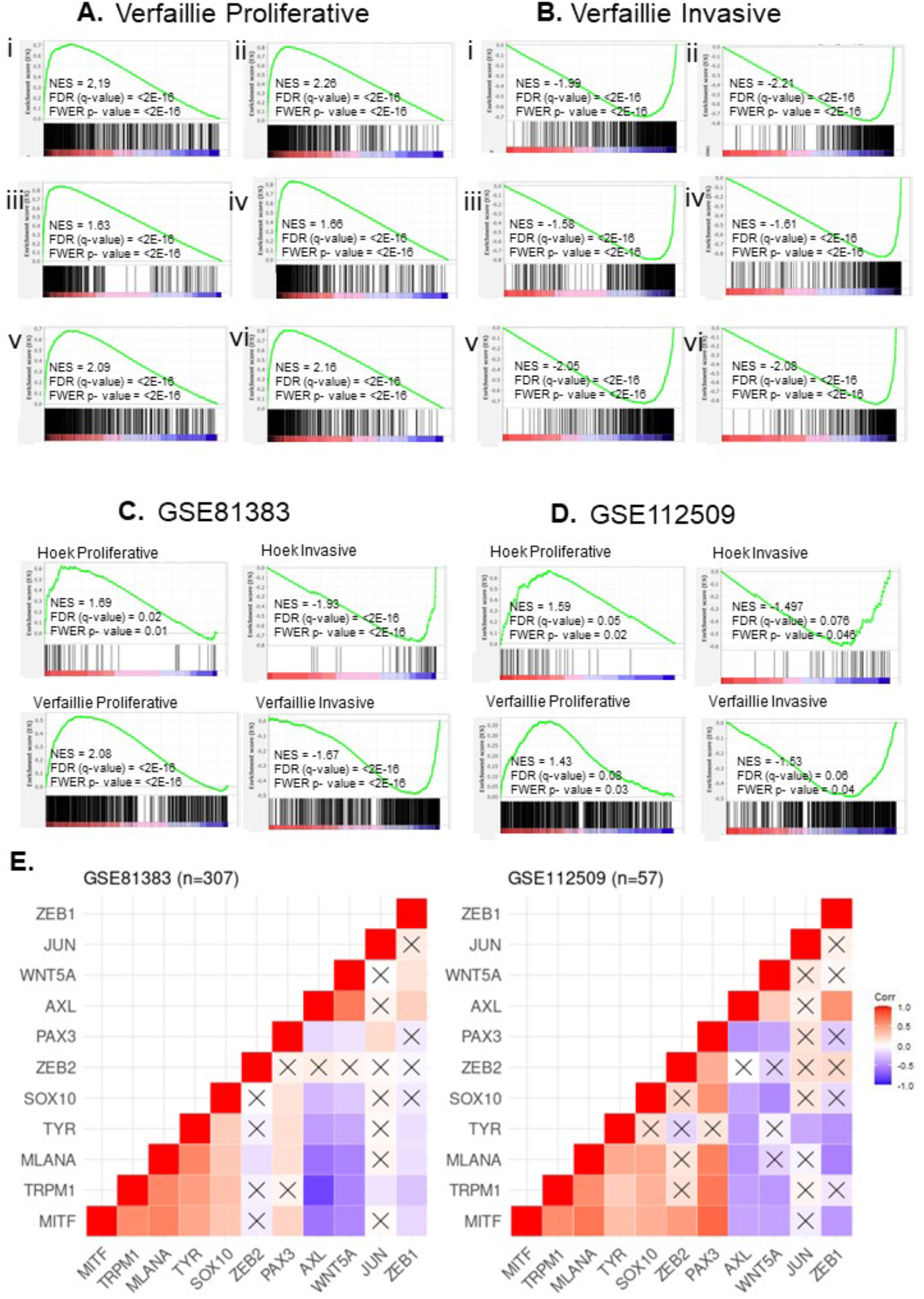
**A,B** GSEA for Verfaillie proliferative and invasive geneset (respectively) for **i**. GSE7127 (n=63) **ii**. GSE80829 (n=53) **iii.** GSE137391 (n=24) **iv.** CCLE (n=59) **v.** GSE10916 (n=50) **vi.** GSE4843 (n=45). GSEA for Hoek Proliferative, Hoek invasive (left to right, top panel), Verfaillie proliferative and Verfaillie invasive geneset (left to right, bottom panel), in GSE81383 (**C.**) and GSE112509 (Primary tumour cells only) (**D.**). **E.** Spearman’s Correlation Coefficient matrix for regulators of phenotypic heterogeneity for GSE81383 and GSE112509 (Primary tumour cells only). Crosses indicate p > 0.05. Colorbar denotes correlation coefficient.

**Fig S3A.**
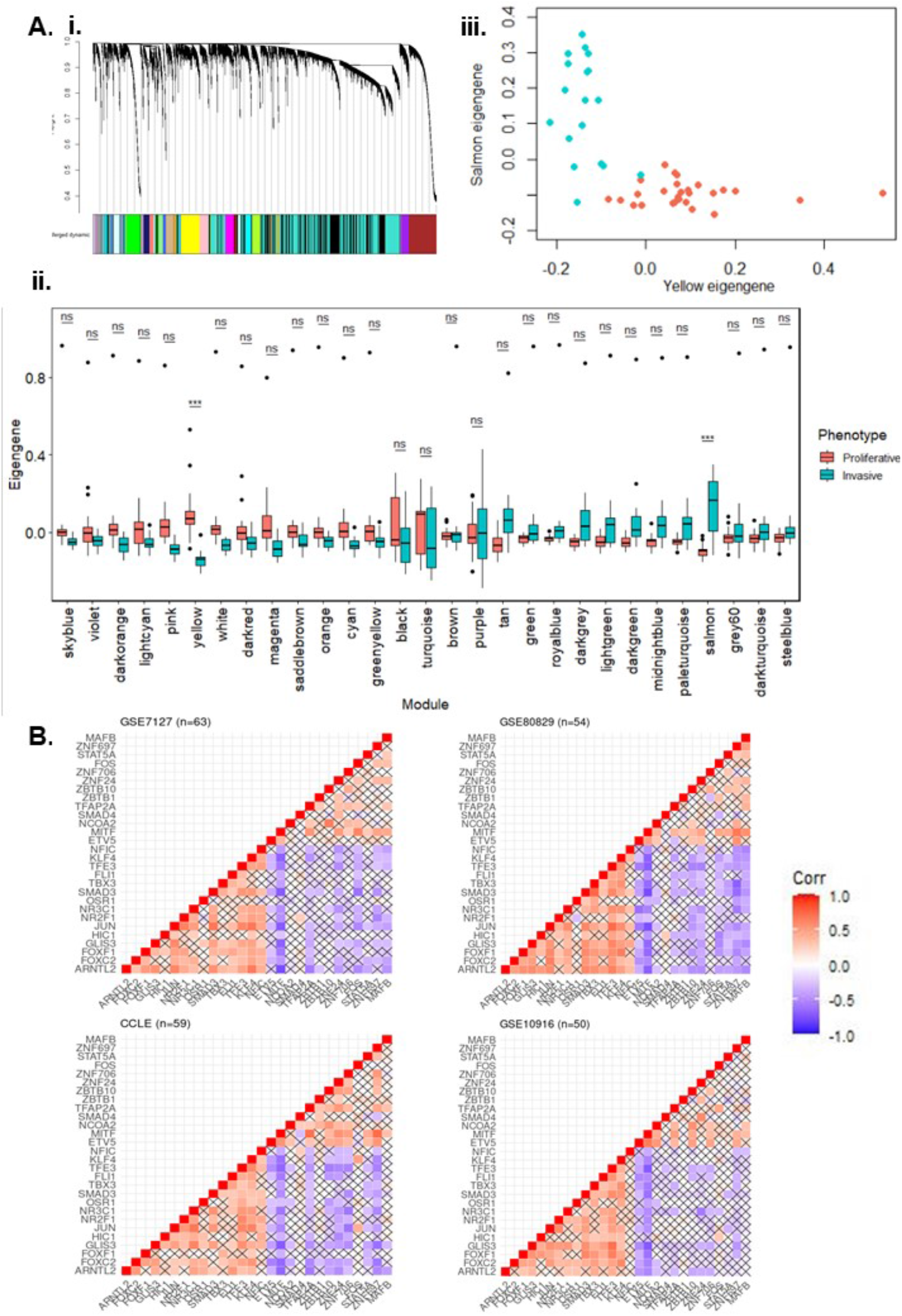
**i.** Dendrogram and modules identified from WGCNA for GE4843 **ii.** Eigengene values for all modules in proliferative and invasive samples. Significance is represented for Bonferonni adjusted p-value using * for p< 0.001, ** for p < 0.0001, *** for p < 0.00001. **iii**. Module eigengene scatterplot for proliferative (Orange) and invasive (cyan) samples for Salmon and yellow modules. **B.** Spearman’s Correlation Coefficient for genes identified as regulators from GSE4843 in CCLE, GSE7127,10916 and GSE80829. Crosses indicate p > 0.05. Colorbars denote correlation coefficient.

**Fig S4.**
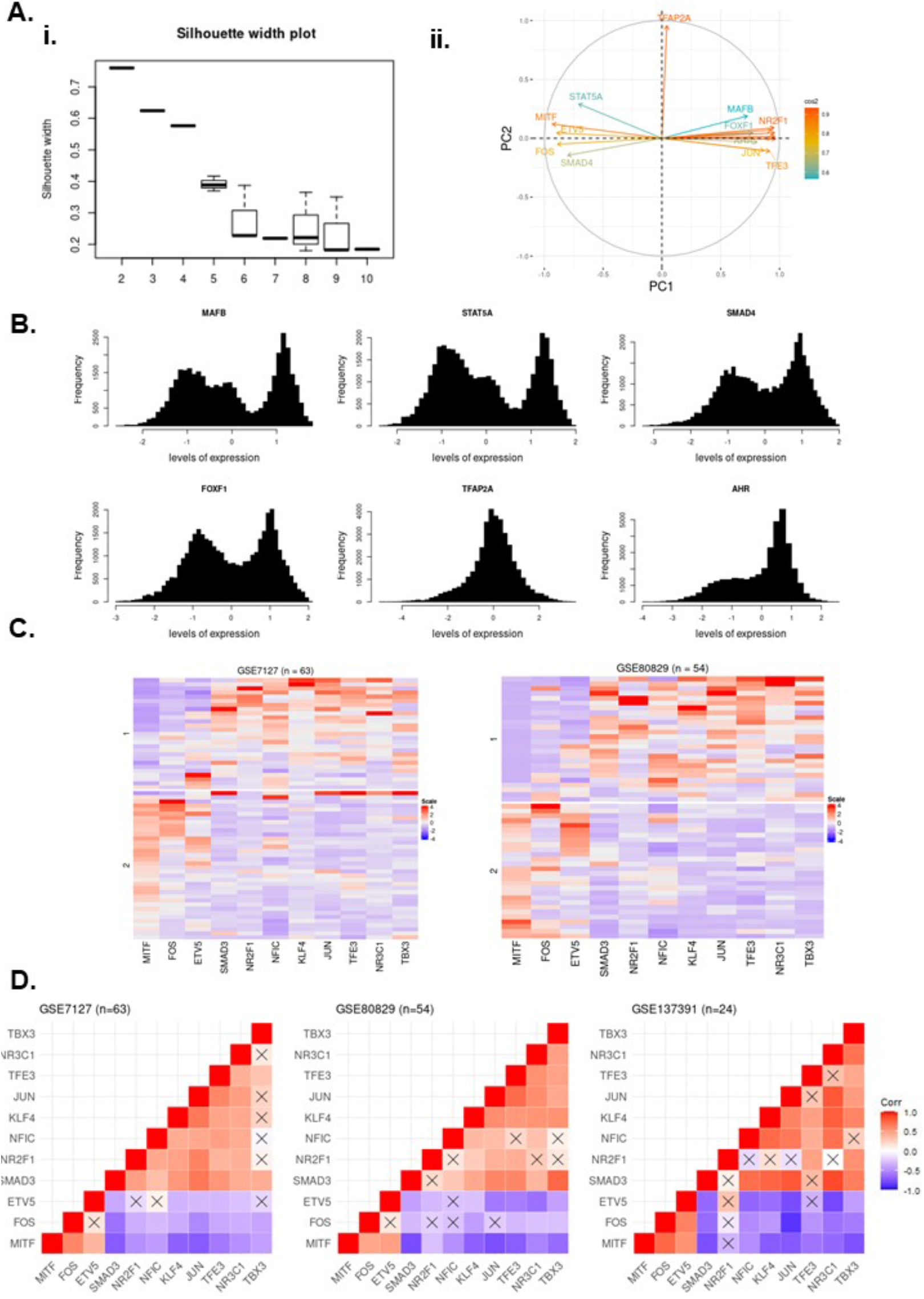
**A. i.** Average silhouette width for k-means clustering measuring for varying number of clusters (k = 2 to 10). Error bars indicate standard deviation. **ii.** Correlation circle with squared cosines of all genes in the network for PC1 and PC2. **B.** Histograms for gene expression simulated using RACIPE. **C.** Differential expression of master regulators in proliferative and invasive samples in GSE7127 and GSE80829 **D.** Spearman’s Correlation Coefficient matrix for a subset of network genes in GSE7127, GSE80829 and GSE137391.

**Fig S5.**
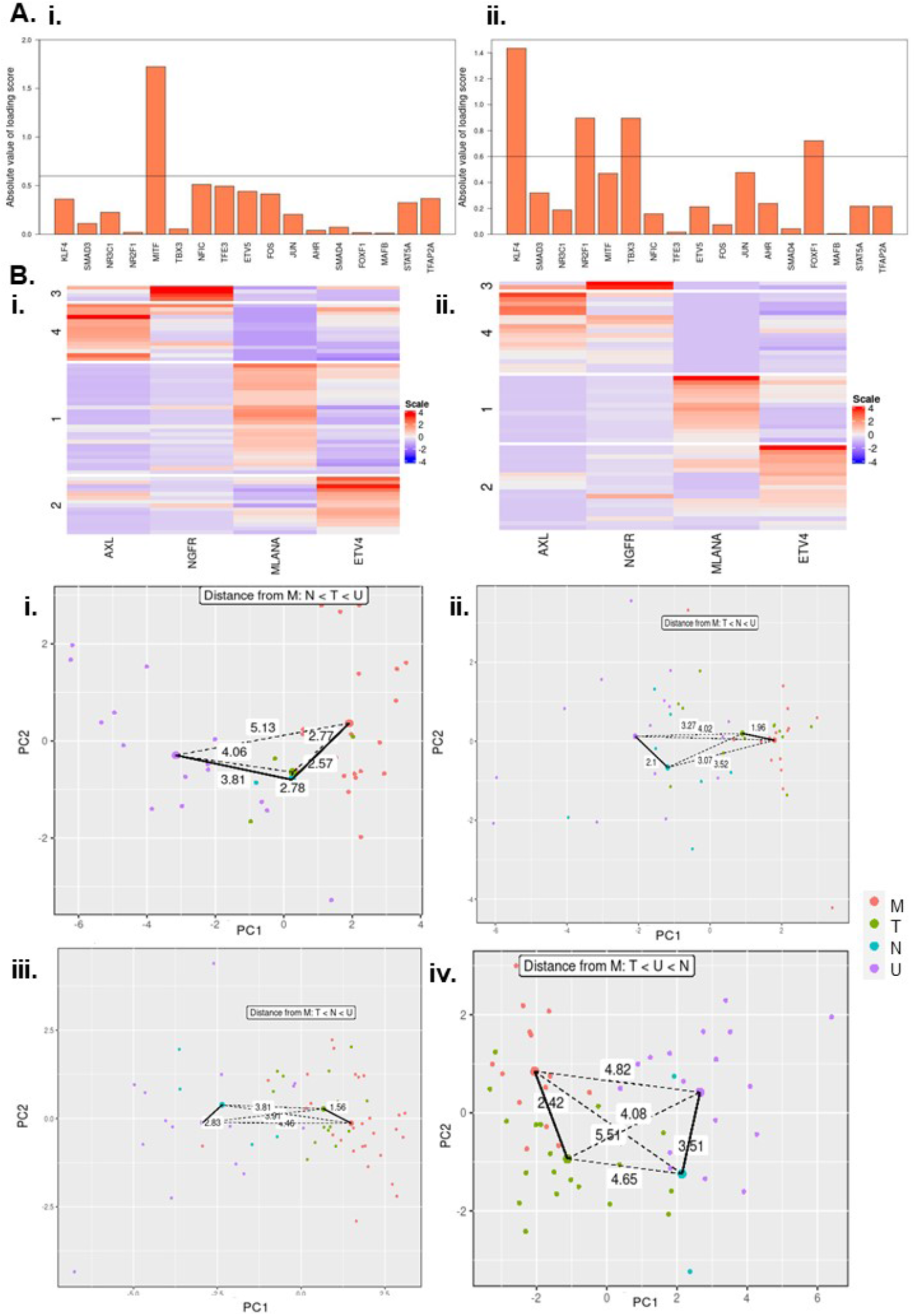
**A**. LDA loading scores for master regulators in **i.** Proliferative subclusters and **ii.** Invasive subclusters. Cut off value of 0.6 was set to select genes. **B.** Heatmap of marker genes expression levels for the 4 phenotypes (AXL, NGFR, MLANA and ETV4) for **i**. GSE7127 **ii**. GSE80829. **C.** All possible trajectories (dotted line), corresponding Euclidean distance and shortest distance to a cluster centre from melanocytic and undifferentiated clusters (solid line) [right] for **i.** GSE4843 **ii**. GSE10916 **iii.** GSE7127 **iv.** GSE80829

**Fig S6.**
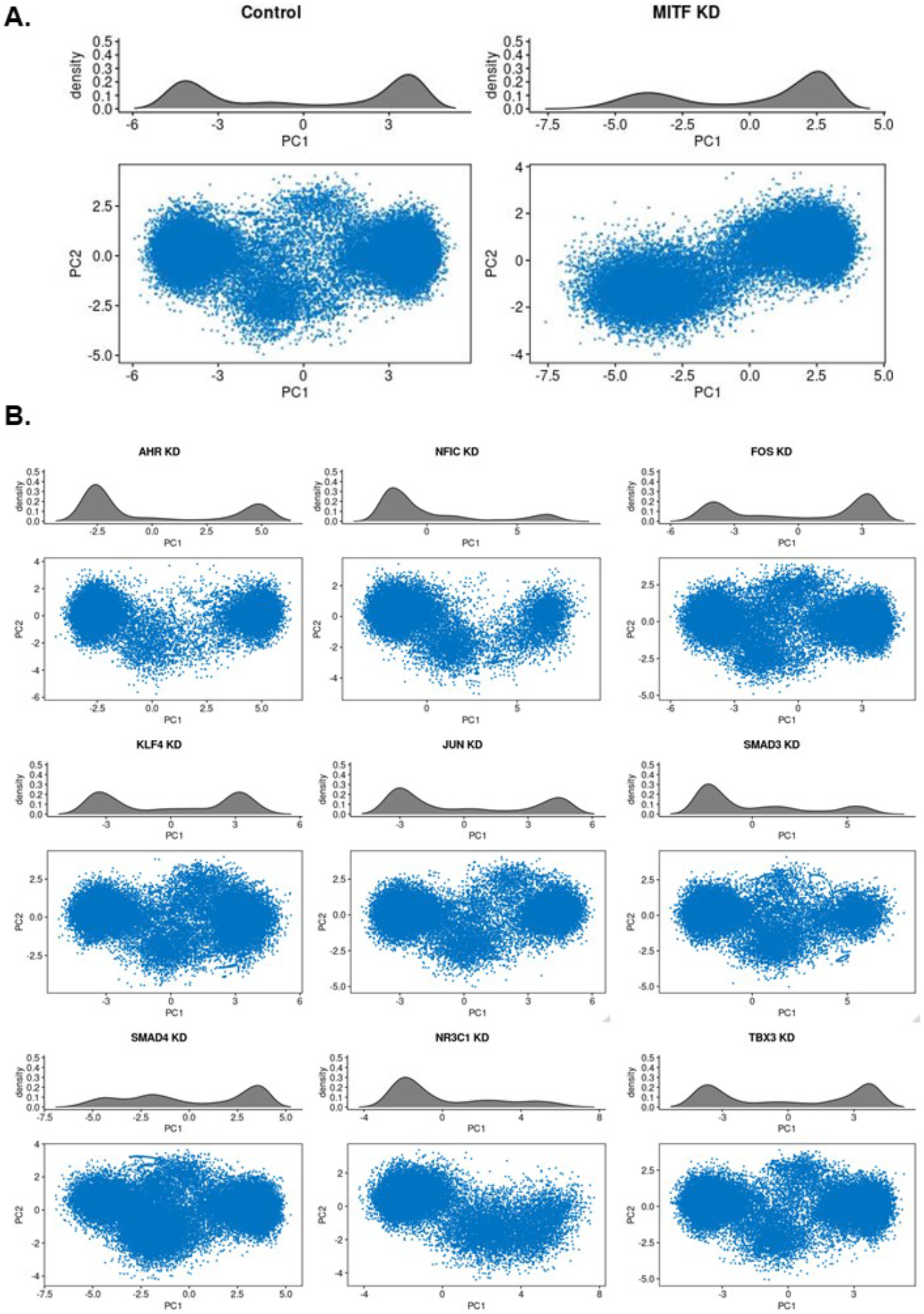
Density distribution of datapoints for **A.** complete network and MITF knock down **B.** Knock-down of rest of the genes in the network

## References

Ahmed, F., and Haass, N.K. (2018). Microenvironment-driven dynamic heterogeneity and phenotypic plasticity as a mechanism of melanoma therapy resistance. Front. Oncol. 8, 173.

Arozarena, I., and Wellbrock, C. (2017). Targeting invasive properties of melanoma cells. FEBS J. 284, 2148–2162.

Augustine, C.K., Toshimitsu, H., Jung, S.H., Zipfel, P.A., Yoo, J.S., Yoshimoto, Y., Selim, M.A., Burchette, J., Beasley, G.M., McMahon, N., et al. (2010). Sorafenib, a multikinase inhibitor, enhances the response of melanoma to regional chemotherapy. Mol. Cancer Ther. 9, 2090–2101.

Aya-Bonilla, C.A., Morici, M., Hong, X., McEvoy, A.C., Sullivan, R.J., Freeman, J., Calapre, L., Khattak, M.A., Meniawy, T., Millward, M., et al. (2020). Detection and prognostic role of heterogeneous populations of melanoma circulating tumour cells. Br. J. Cancer 122, 1059–1067.

Barretina, J., Caponigro, G., Stransky, N., Venkatesan, K., Margolin, A. a., Kim, S., Wilson, C.J., Lehár, J., Kryukov, G. V., Sonkin, D., et al. (2012). The Cancer Cell Line Encyclopedia enables predictive modelling of anticancer drug sensitivity. Nature 483, 603–607.

Bettum, I.J., Gorad, S.S., Barkovskaya, A., Pettersen, S., Moestue, S.A., Vasiliauskaite, K., Tenstad, E., Øyjord, T., Risa, Ø., Nygaard, V., et al. (2015). Metabolic reprogramming supports the invasive phenotype in malignant melanoma. Cancer Lett. 366, 71–83.

Bocci, F., Mandal, S., Tejaswi, T., and Jolly, M.K. (2021). Investigating epithelial-mesenchymal heterogeneity of tumors and circulating tumor cells with transcriptomic analysis and biophysical modeling. Comput. Syst. Oncol. in press.

Boshuizen, J., Koopman, L.A., Krijgsman, O., Shahrabi, A., Van Den Heuvel, E.G., Ligtenberg, M.A., Vredevoogd, D.W., Kemper, K., Kuilman, T., Song, J.Y., et al. (2018). Cooperative targeting of melanoma heterogeneity with an AXL antibody-drug conjugate and BRAF/MEK inhibitors. Nat. Med. 24, 203–212.

Caramel, J., Papadogeorgakis, E., Hill, L., Browne, G.J., Richard, G., Wierinckx, A., Saldanha, G., Sborne, J., Hutchinson, P., Tse, G., et al. (2013). A Switch in the Expression of Embryonic EMT-Inducers Drives the Development of Malignant Melanoma. Cancer Cell 24, 466–480.

Chauhan, L., Ram, U., Hari, K., and Jolly, M.K. (2020). Topological signatures in regulatory network enable phenotypic heterogeneity in small cell lung cancer. BioRxiv 362228.

Chen, E.Y., Tan, C.M., Kou, Y., Duan, Q., Wang, Z., Meirelles, G. V., Clark, N.R., and Ma’ayan, A. (2013). Enrichr: Interactive and collaborative HTML5 gene list enrichment analysis tool. BMC Bioinformatics 14, 128.

Denecker, G., Vandamme, N., Akay, Ö., Koludrovic, D., Taminau, J., Lemeire, K., Gheldof, A., De Craene, B., Van Gele, M., Brochez, L., et al. (2014). Identification of a ZEB2-MITF-ZEB1 transcriptional network that controls melanogenesis and melanoma progression. Cell Death Differ. 21, 1250–1261.

Deng, Y., Chakraborty, P., Jolly, M.K., and Levine, H. (2020). A theoretical approach to coupling the epithelial-mesenchymal transition (EMT) to extracellular matrix (ECM) stiffness via LOXL2. BioRxiv 429859.

Fallahi-Sichani, M., Becker, V., Izar, B., Baker, G.J., Lin, J.-R., Boswell, S.A., Shah, P., Rotem, A., Garraway, L.A., and Sorger, P.K. (2017). Adaptive resistance of melanoma cells to RAF inhibition via reversible induction of a slowly dividing de-differentiated state. Mol Syst Biol 13, 905.

Falletta, P., Sanchez-del-Campo, L., Chauhan, J., Effern, M., Kenyon, A., Kershaw, C.J., Siddaway, R., Lisle, R., Freter, R., Daniels, M.J., et al. (2017). Translation reprogramming is an evolutionarily conserved driver of phenotypic plasticity and therapeutic resistance in melanoma. Genes Dev. 31, 18–33.

Fane, M.E., Chhabra, Y., Smith, A.G., and Sturm, R.A. (2019). BRN2, a POUerful driver of melanoma phenotype switching and metastasis. Pigment Cell Melanoma Res. 32, 9–24.

Feldker, N., Ferrazzi, F., Schuhwerk, H., Widholz, S.A., Guenther, K., Frisch, I., Jakob, K., Kleemann, J., Riegel, D., Bönisch, U., et al. (2020). Genome-wide cooperation of EMT transcription factor ZEB 1 with YAP and AP −1 in breast cancer. EMBO J. 39, e103209.

Floratos, A., Smith, K., Ji, Z., Watkinson, J., and Califano, A. (2010). geWorkbench: An open source platform for integrative genomics. Bioinformatics 26, 1779–1780.

Gerber, T., Willscher, E., Loeffler-Wirth, H., Hopp, L., Schadendorf, D., Schartl, M., Anderegg, U., Camp, G., Treutlein, B., Binder, H., et al. (2017). Mapping heterogeneity in patient-derived melanoma cultures by single-cell RNA-seq. Oncotarget 8, 846–862.

Goetz, H., Melendez-Alvarez, J.R., Chen, L., and Tian, X.-J. (2020). A plausible accelerating function of intermediate states in cancer metastasis. PLOS Comput. Biol. 16, e1007682.

Goldman, A., Majumder, B., Dhawan, A., Ravi, S., Goldman, D., Kohandel, M., Majumder, P.K., and Sengupta, S. (2015). Temporally sequenced anticancer drugs overcome adaptive resistance by targeting a vulnerable chemotherapy-induced phenotypic transition. Nat. Commun. 6, 6139.

Hartigan, J.A., and Hartigan, P.M. (1985). The Dip Test of Unimodality. Ann. Stat. 13, 70–84.

Hoek, K.S., Schlegel, N.C., Brafford, P., Sucker, A., Ugurel, S., Kumar, R., Weber, B.L., Nathanson, K.L., Phillips, D.J., Herlyn, M., et al. (2006). Metastatic potential of melanomas defined by specific gene expression profiles with no BRAF signature. Pigment Cell Res. 19, 290–302.

Hoek, K.S., Eichhoff, O.M., Schlegel, N.C., Döbbeling, U., Kobert, N., Schaerer, L., Hemmi, S., and Dummer, R. (2008). In vivo switching of human melanoma cells between proliferative and invasive states. Cancer Res. 68, 650–656.

Huang, B., Lu, M., Jia, D., Ben-Jacob, E., Levine, H., and Onuchic, J.N. (2017). Interrogating the topological robustness of gene regulatory circuits by randomization. PLoS Comput. Biol. 13, e1005456.

Hugo, W., Shi, H., Sun, L., Piva, M., Song, C., Kong, X., Moriceau, G., Hong, A., Dahlman, K.B., Johnson, D.B., et al. (2015). Non-genomic and Immune Evolution of Melanoma Acquiring MAPKi Resistance. Cell 162, 1271–1285.

Jia, D., Paudel, B.B., Hayford, C.E., Hardeman, K.N., Levine, H., Onuchic, J.N., and Quaranta, V. (2020a). Drug-Tolerant Idling Melanoma Cells Exhibit Theory-Predicted Metabolic Low-Low Phenotype. Front. Oncol. 10, 1426.

Jia, W., Deshmukh, A., Mani, S.A., Jolly, M.K., and Levine, H. (2019). A possible role for epigenetic feedback regulation in the dynamics of the Epithelial-Mesenchymal Transition (EMT). Phys. Biol. 16, 066004.

Jia, W., Tripathi, S., Chakraborty, P., Chedere, A., Rangarajan, A., Levine, H., and Jolly, M.K. (2020b). Epigenetic feedback and stochastic partitioning during cell division can drive resistance to EMT. Oncotarget 11, 2611–2624.

Johansson, P., Pavey, S., and Hayward, N. (2007). Confirmation of a BRAF mutation-associated gene expression signature in melanoma. Pigment Cell Res. 20, 216–221.

Jolly, M.K., and Celia-Terrassa, T. (2019). Dynamics of Phenotypic Heterogeneity Associated with EMT and Stemness during Cancer Progression. J. Clin. Med. 8, 1542.

Karacosta, L.G., Anchang, B., Ignatiadis, N., Kimmey, S.C., Benson, J.A., Shrager, J.B., Tibshirani, R., Bendall, S.C., and Plevritis, S.K. (2019). Mapping Lung Cancer Epithelial-Mesenchymal Transition States and Trajectories with Single-Cell Resolution. Nat. Commun. 10, 5587.

Karki, P., Angardi, V., Mier, J.C., and Orman, M.A. (2021). A Transient Metabolic State In Melanoma Persister Cells Mediated By Chemotherapeutic Treatments. BioRxiv 432154.

Kunz, M., Löffler-Wirth, H., Dannemann, M., Willscher, E., Doose, G., Kelso, J., Kottek, T., Nickel, B., Hopp, L., Landsberg, J., et al. (2018). RNA-seq analysis identifies different transcriptomic types and developmental trajectories of primary melanomas. Oncogene 37, 6136–6151.

Lambert, S.A., Jolma, A., Campitelli, L.F., Das, P.K., Yin, Y., Albu, M., Chen, X., Taipale, J., Hughes, T.R., and Weirauch, M.T. (2018). The Human Transcription Factors. Cell 172, 650–655.

Langfelder, P., and Horvath, S. (2008). WGCNA: an R package for weighted correlation network analysis. BMC Bioinformatics 9, 559.

Li, C., Hong, T., and Nie, Q. (2016). Quantifying the landscape and kinetic paths for epithelial– mesenchymal transition from a core circuit. Phys. Chem. Chem. Phys. 18, 17949–17956.

Liguoro, D., Fattore, L., Mancini, R., and Ciliberto, G. (2020). Drug tolerance to target therapy in melanoma revealed at single cell level: What next? Biochim. Biophys. Acta - Rev. Cancer 1874, 188440.

Lionetti, M.C., Cola, F., Chepizhko, O., Fumagalli, M.R., Font-Clos, F., Ravasio, R., Minucci, S., Canzano, P., Camera, M., Tiana, G., et al. (2020). MicroRNA-222 Regulates Melanoma Plasticity. J. Clin. Med. 9, 2573.

Luo, M., Shang, L., Brooks, M.D., Jiagge, E., Zhu, Y., Buschhaus, J.M., Conley, S., Fath, M.A., Davis, A., Gheordunescu, E., et al. (2018). Targeting Breast Cancer Stem Cell State Equilibrium through Modulation of Redox Signaling. Cell Metab. 28, 69–86.

Margolin, A.A., Nemenman, I., Basso, K., Wiggins, C., Stolovitzky, G., Dalla Favera, R., and Califano, A. (2006). ARACNE: an algorithm for the reconstruction of gene regulatory networks in a mammalian cellular context. BMC Bioinformatics 7 Suppl 1, S7.

Müller, J., Krijgsman, O., Tsoi, J., Robert, L., Hugo, W., Song, C., Kong, X., Possik, P.A., Cornelissen-Steijger, P.D.M., Foppen, M.H.G., et al. (2014). Low MITF/AXL ratio predicts early resistance to multiple targeted drugs in melanoma. Nat. Commun. 5, 5712.

Pastushenko, I., Brisebarre, A., Sifrim, A., Fioramonti, M., Revenco, T., Boumahdi, S., Van Keymeulen, A., Brown, D., Moers, V., Lemaire, S., et al. (2018). Identification of the tumour transition states occurring during EMT. Nature 556, 463–468.

Paudel, B.B., Harris, L.A., Hardeman, K.N., Abugable, A.A., Hayford, C.E., Tyson, D.R., and Quaranta, V. (2018). A Nonquiescent “Idling” Population State in Drug-Treated, BRAF-Mutated Melanoma. Biophys. J. 114, 1499–1511.

Pfister, R., Schwarz, K.A., Janczyk, M., Dale, R., and Freeman, J.B. (2013). Good things peak in pairs: A note on the bimodality coefficient. Front. Psychol. 4, 700.

Rambow, F., Rogiers, A., Marin-Bejar, O., Aibar, S., Femel, J., Dewaele, M., Karras, P., Brown, D., Chang, Y.H., Debiec-Rychter, M., et al. (2018). Toward Minimal Residual Disease-Directed Therapy in Melanoma. Cell 174, 843–855.e59.

Rambow, F., Marine, J.C., and Goding, C.R. (2019). Melanoma plasticity and phenotypic diversity: Therapeutic barriers and opportunities. Genes Dev. 33, 1295–1318.

Rebecca, V.W., and Herlyn, M. (2020). Nongenetic Mechanisms of Drug Resistance in Melanoma. Annu. Rev. Cancer Biol. 4, 315–330.

Reinhardt, J., Landsberg, J., Schmid-Burgk, J.L., Ramis, B.B., Bald, T., Glodde, N., Lopez-Ramos, D., Young, A., Ngiow, S.F., Nettersheim, D., et al. (2017). MAPK signaling and inflammation link melanoma phenotype switching to induction of CD73 during immunotherapy. Cancer Res. 77, 4697–4709.

Riesenberg, S., Groetchen, A., Siddaway, R., Bald, T., Reinhardt, J., Smorra, D., Kohlmeyer, J., Renn, M., Phung, B., Aymans, P., et al. (2015). MITF and c-Jun antagonism interconnects melanoma dedifferentiation with pro-inflammatory cytokine responsiveness and myeloid cell recruitment. Nat. Commun. 6, 8755.

Roesch, A., Fukunaga-Kalabis, M., Schmidt, E.C., Zabierowski, S.E., Brafford, P.A., Vultur, A., Basu, D., Gimotty, P., Vogt, T., and Herlyn, M. (2010). A Temporarily Distinct Subpopulation of Slow-Cycling Melanoma Cells Is Required for Continuous Tumor Growth. Cell 141, 583–594.

Rowling, E.J., Miskolczi, Z., Nagaraju, R., Wilcock, D.J., Wang, P., Telfer, B., Li, Y., Lasheras-Otero, I., Redondo-Muñoz, M., Sharrocks, A.D., et al. (2020). Cooperative behaviour and phenotype plasticity evolve during melanoma progression. Pigment Cell Melanoma Res. 33, 695–708.

Schuh, L., Saint-Antoine, M., Sanford, E.M., Emert, B.L., Singh, A., Marr, C., Raj, A., and Goyal, Y. (2020). Gene Networks with Transcriptional Bursting Recapitulate Rare Transient Coordinated High Expression States in Cancer. Cell Syst. 10, P363–378.E12.

Shaffer, S.M., Dunagin, M.C., Torborg, S.R., Torre, E.A., Emert, B., Krepler, C., Beqiri, M., Sproesser, K., Brafford, P.A., Xiao, M., et al. (2017). Rare cell variability and drug-induced reprogramming as a mode of cancer drug resistance. Nature 546, 431–435.

Smith, M.P., Rana, S., Ferguson, J., Rowling, E.J., Flaherty, K.T., Wargo, J.A., Marais, R., and Wellbrock, C. (2019). A PAX3/BRN2 rheostat controls the dynamics of BRAF mediated MITF regulation in MITFhigh/AXLlow melanoma. Pigment Cell Melanoma Res. 32, 280–291.

Su, Y., Wei, W., Robert, L., Xue, M., Tsoi, J., Garcia-diaz, A., Homet, B.M., Kim, J., Ng, R.H., Lee, J.W., et al. (2017). Single-cell analysis resolves the cell state transition and signaling dynamics associated with melanoma drug-induced resistance. Proc Natl Acad Sci U S A 114, 13679–13684.

Su, Y., Bintz, M., Yang, Y., Robert, L., Ng, A.H.C., Liud, V., Ribas, A., Heath, J.R., and Wei, W. (2019). Phenotypic heterogeneity and evolution of melanoma cells associated with targeted therapy resistance. PLoS Comput. Biol. 15, e1007034.

Subramanian, A., Tamayo, P., Mootha, V.K., Mukherjee, S., Ebert, B.L., Gillette, M.A., Paulovich, A., Pomeroy, S.L., Golub, T.R., Lander, E.S., et al. (2005). Gene set enrichment analysis: A knowledge-based approach for interpreting genome-wide expression profiles. Proc Natl Acad Sci U S A 102, 15545–15550.

Sullivan, R.J., and Flaherty, K.T. (2013). Resistance to BRAF-targeted therapy in melanoma. Eur. J. Cancer 49, 1297–1304.

Sun, C., Wang, L., Huang, S., Heynen, G.J.J.E., Prahallad, A., Robert, C., Haanen, J., Blank, C., Wesseling, J., Willems, S.M., et al. (2014). Reversible and adaptive resistance to BRAF (V600E) inhibition in melanoma. Nature 508, 118–122.

Tirosh, I., Izar, B., Prakadan, S.M., Ii, M.H.W., Treacy, D., Trombetta, J.J., Rotem, A., Rodman, C., Lian, C., Murphy, G., et al. (2016). Dissecting the multicellular ecosystem of metastatic melanoma by single-cell RNA-seq. Science 352, 189–196.

Tremblay, B.L., Guénard, F., Lamarche, B., Pérusse, L., and Vohl, M.C. (2019). Weighted gene co-expression network analysis to explain the relationship between plasma total carotenoids and lipid profile. Genes Nutr. 14, 16.

Tripathi, S., Chakraborty, P., Levine, H., and Jolly, M.K. (2020). A mechanism for epithelial-mesenchymal heterogeneity in a population of cancer cells. PLoS Comput Biol 16, e1007619.

Tsoi, J., Robert, L., Paraiso, K., Galvan, C., Sheu, K.M., Lay, J., Wong, D.J.L., Atefi, M., Shirazi, R., Wang, X., et al. (2018). Multi-stage Differentiation Defines Melanoma Subtypes with Differential Vulnerability to Drug-Induced Iron-Dependent Oxidative Stress. Cancer Cell 33, 890–904.e5.

Udyavar, A.A., Wooten, D.J., Hoeksema, M., Bansal, M., Califano, M., Estrada, L., Schnell, S., Irish, J.M., Massion, P.P., and Quaranta, V. (2017). Novel Hybrid Phenotype Revealed in Small Cell Lung Cancer by a Transcription Factor Network Model That Can Explain Tumor Heterogeneity. Cancer Res. 77, 1063–1074.

Vandamme, N., Denecker, G., Bruneel, K., Blancke, G., Akay, O., Taminau, J., de Coninck, J., de Smedt, E., Skrypek, N., van Loocke, W., et al. (2020). The EMT transcription factor ZEB2 promotes proliferation of primary and metastatic melanoma while suppressing an invasive, mesenchymal-like phenotype. Cancer Res. 80, 2983–2995.

Vazquez, F., Lim, J.H., Chim, H., Bhalla, K., Girnun, G., Pierce, K., Clish, C.B., Granter, S.R., Widlund, H.R., Spiegelman, B.M., et al. (2013). PGC1α Expression Defines a Subset of Human Melanoma Tumors with Increased Mitochondrial Capacity and Resistance to Oxidative Stress. Cancer Cell 23, 287–301.

Verfaillie, A., Imrichova, H., Atak, Z.K., Dewaele, M., Rambow, F., Hulselmans, G., Christiaens, V., Svetlichnyy, D., Luciani, F., Van Den Mooter, L., et al. (2015). Decoding the regulatory landscape of melanoma reveals TEADS as regulators of the invasive cell state. Nat. Commun. 6, 6683.

Vivas-García, Y., Falletta, P., Liebing, J., Louphrasitthiphol, P., Feng, Y., Chauhan, J., Scott, D.A., Glodde, N., Chocarro-Calvo, A., Bonham, S., et al. (2020). Lineage-Restricted Regulation of SCD and Fatty Acid Saturation by MITF Controls Melanoma Phenotypic Plasticity. Mol. Cell 77, 120–137.e9.

Weeraratna, A.T., Jiang, Y., Hostetter, G., Rosenblatt, K., Duray, P., Bittner, M., and Trent, J.M. (2002). Wnt5a signaling directly affects cell motility and invasion of metastatic melanoma. Cancer Cell 1, 279–288.

Wei, S.C., Fattet, L., Tsai, J.H., Guo, Y., Pai, V.H., Majeski, H.E., Chen, A.C., Sah, R.L., Taylor, S.S., Engler, A.J., et al. (2015). Matrix stiffness drives epithelial–mesenchymal transition and tumour metastasis through a TWIST1–G3BP2 mechanotransduction pathway. Nat. Cell Biol. 17, 678–688.

Wouters, J., Kalender-Atak, Z., Minnoye, L., Spanier, K.I., De Waegeneer, M., Bravo González-Blas, C., Mauduit, D., Davie, K., Hulselmans, G., Najem, A., et al. (2020). Robust gene expression programs underlie recurrent cell states and phenotype switching in melanoma. Nat. Cell Biol. 22, 986–998.

Yu, M., Bardia, A., Wittner, B.S., Stott, S.L., Smas, M.E., Ting, D.T., Isakoff, S.J., Ciciliano, J.C., Wells, M.N., Shah, A.M., et al. (2013). Circulating breast tumor cells exhibit dynamic changes in epithelial and mesenchymal composition. Science 339, 580–584.

Zhou, J.X., and Huang, S. (2011). Understanding gene circuits at cell-fate branch points for rational cell reprogramming. Trends Genet. 27, 55–62.

